# Complete absence of GLUT1 does not impair human terminal erythroid differentiation

**DOI:** 10.1101/2024.01.10.574621

**Authors:** CM Freire, NR King, M Dzieciatkowska, D Stephenson, PL Moura, J.G.G Dobbe, GJ Streekstra, A D’Alessandro, AM Toye, TJ Satchwell

## Abstract

The Glucose transporter 1 (GLUT1) is one of the most abundant proteins within the erythrocyte membrane and is required for glucose and dehydroascorbic acid (Vitamin C precursor) transport. It is widely recognized as a key protein for red cell structure, function, and metabolism. Previous reports highlighted the importance of GLUT1 activity within these uniquely glycolysis-dependent cells, in particular for increasing antioxidant capacity needed to avoid irreversible damage from oxidative stress in humans. However, studies of glucose transporter roles in erythroid cells are complicated by species-specific differences between humans and mice. Here, using CRISPR-mediated gene editing of immortalized erythroblasts and adult CD34+ hematopoietic progenitor cells, we generate committed human erythroid cells completely deficient in expression of GLUT1. We show that absence of GLUT1 does not impede human erythroblast proliferation, differentiation, or enucleation. This work demonstrates for the first-time generation of enucleated human reticulocytes lacking GLUT1. The GLUT1-deficient reticulocytes possess no tangible alterations to membrane composition or deformability in reticulocytes. Metabolomic analyses of GLUT1-deficient reticulocytes reveal hallmarks of reduced glucose import, downregulated metabolic processes and upregulated AMPK-signalling, alongside alterations in antioxidant metabolism, resulting in increased osmotic fragility and metabolic shifts indicative of higher oxidant stress. Despite detectable metabolic changes in GLUT1 deficient reticulocytes, the absence of developmental phenotype, detectable proteomic compensation or impaired deformability comprehensively alters our understanding of the role of GLUT1 in red blood cell structure, function and metabolism. It also provides cell biological evidence supporting clinical consensus that reduced GLUT1 expression does not cause anaemia in GLUT1 deficiency syndrome.

**Key Points:**

- GLUT1 knockout does not affect erythroid differentiation and minimally impacts reticulocyte membrane composition
- Metabolic adaptation facilitates reticulocyte tolerance of GLUT1 absence

## Introduction

Glucose is an essential source of energy, sustaining life across multiple organisms. Glucose transport in vertebrates is assured by the SLC2 (GLUT) family, with GLUT1, encoded by the SLC2A1 gene, as the _rst characterised glucose transporter^1^. Of all the cell lineages within the human body, human erythrocytes express the highest levels of the GLUT1 transporter, with approximately 200,000 copies per cell accounting for 10% of the total membrane protein mass of the erythrocyte^2^.

Within the red blood cell (RBC) membrane, GLUT1 is located in the junctional multiprotein complex, where it associates with the underlying spectrin cytoskeleton via cytoskeletal adaptor proteins such as dematin and adducin^3,4^. GLUT1 also interacts with stomatin, a membrane associated protein that binds to cholesterol and is involved in membrane scaffolding,^5^ and GLUT1 1 is proposed to associate with band 3 (Anion exchanger 1) via its C-terminal domain.

GLUT1 transports glucose into erythroid cells, providing a source of energy for ATP generation either through the Krebs cycle during erythropoiesis or via maintaining glycolysis in the mature RBC following the loss of mitochondria that occurs during terminal differentiation^6^. Characteristic features of RBC such as their extensive capacity for deformation during capillary transit, influenced by ATP dependent phosphorylation, calcium, and cell volume homeostasis, are intrinsically linked to the metabolic status of the erythrocyte, and therefore governed by the availability of glucose^7,8^. Somewhat paradoxically, whilst RBC are exclusively reliant upon anaerobic glycolysis to meet their energetic requirements, glucose transport decreases during erythropoiesis (despite increased expression of GLUT1), due to a shift to transport of the vitamin C precursor dehydroascorbic acid (DHA) favoured in humans and other primates that lack the ability to synthesise the potent antioxidant ascorbic acid (Vitamin C)^9^. The alteration in GLUT1 transport substrate selectivity is driven by the association of stomatin^9^.

The complete absence of GLUT1 in humans has not been reported and is presumed to be embryonically lethal. GLUT1 deficiency syndrome (G1DS) is a neurodevelopmental disorder that results from haploinsuffiency for GLUT1 due to specific mutations in the SLC2A1 gene. This leads to impaired glucose transport into the brain and RBCs^10^ manifesting as a range of neurological symptoms, including epilepsy, developmental delay, and movement disorders^11–13^. The neurological disease can be modelled in mice by introducing heterozygous GLUT1 mutations^14^, but since GLUT1 is only expressed in erythrocytes during the murine perinatal period and is rapidly replaced by GLUT4 in adult mouse erythrocytes, the impact of haploinsufficiency of GLUT1 on RBC cannot be fully modelled in mice. Interestingly, despite the perceived pivotal role of GLUT1 in RBC function, patients with G1DS typically do not exhibit any haematological phenotypes arising from the deficiency of this protein^15^. The exact reason for this “lack of RBC phenotype” remains unclear and represents an intriguing area of investigation.

In this study we exploit CRISPR-mediated gene editing of both immortalised erythroblasts and adult haematopoietic stem cells, generating novel GLUT1 KO erythroid cells to establish the impact of total loss of GLUT1 on RBC formation, membrane protein composition and stability, and metabolism. Surprisingly, we show that absence of GLUT1 is well tolerated by differentiating erythroblasts with no impairment of proliferation, enucleation, or tangible alterations to resulting reticulocyte membrane composition or deformability, challenging prevailing dogma surrounding the necessity of this abundant protein for RBC development, structure, and function.

## Materials and Methods

### Source Material

All human blood source material was provided with written informed consent for research use given in accordance with the Declaration of Helsinki (NHSBT, Filton, Bristol). The research into the mechanisms of erythropoiesis was reviewed and approved by the Bristol Research Ethics committee (REC Number 12/SW/0199).

### Antibodies

See *Supplementary Methods* for detailed list of used antibodies and reagents.

### CD34+ and BEL-A culture

CD34+ cells were isolated, expanded and differentiated as previously described^16^ and further detailed in *Supplemental Methods*.

BEL-A (Bristol Erythroid Line – Adult) cells were cultured as previously described^17,18^ For expansion, cells were seeded at 0.5-1×10^5^ cells/ml in expansion medium, consisting of StemSpan SFEM (Stem Cell Technologies) supplemented with 50ng/ml Stem cell Factor (SCF, R&D Systems), 3U/ml EPO (Neocormon), 1μM dexamethasone (Sigma-Aldrich) and 1μg/ml doxycycline (Clontech). Cells were incubated at 37°C, 5% CO2, with complete medium changes every 48-72h. BEL-A differentiation protocol was performed as described in King et al^19^,. Briefly, cells were seeded at 1.5×10^5^/ml in primary differentiation medium supplemented with 1ng/ml interleukin-3 (IL-3, R&D Systems), 10ng/ml SCF and 1μg/ml doxycycline. After 2 days, cells were reseeded at 3×10^5^/ml in fresh medium. On day 4, cells were reseeded at 5×10^5^/ml in fresh medium without doxycycline. On day 7, cells were transferred to tertiary medium (no SCF, IL-3 or doxycycline) at 1×10^6^/ml. Another complete media change was performed on day 9. Cells were analysed on day 10 or 11.

### CRISPR-editing of cells

BEL-A cells were nucleofected using a 4D-NucleofectorTM System with 20μL Nucleocuvette Strip (Lonza) in combination with the P3 Primary Cell Buffer Kit (Lonza). Per sample, 2×10^5^ cells were transfected with preincubated 18pmol of Cas9 (TrueCut Cas9 Protein v2, ThermoFisher) and 45pmol of either SLC2A1 targeting sgRNA (5’-GGATGCTCTCCCCATAGCGG, Synthego) or non-targeting (NT) control (Negative Control Scrambled sgRNA#1, Synthego), using program DZ-100. For CD34^+^ isolated cells, nucleofection was performed on day 3 after isolation, where 5×10^5^ cells were nucleofected with 50pmol Cas9 and 125pmol sgRNA using electroporation program EO-100.

### Flow cytometry and Fluorescence-activated Cell Sorting (FACS)

For flow cytometry, cells were fixed (1% paraformaldehyde, 0.0075% glutaraldehyde) to prevent antibody agglutination, except for assays using GLUT1.RBD. Samples were labelled with primary antibodies in PBSAG (PBS with 1% (w/v) Glucose and 0.5%(w/v) |Bovine Serum Albumin (BSA)) supplemented with extra 1%(w/v) BSA for 30min, in the dark. METAFORA’s GLUT1.RBD was incubated at 37°C and all remaining antibodies incubated at 4°C. Cells were washed 2x PBSAG and, if required, incubated with appropriate secondary antibody under the same conditions as described for primary. Cells were washed 2x with PBSAG and analysed on a Miltenyi MACSQuant 10 flow cytometer. Data was analysed using FlowJo v10.7 (FlowJo). Reticulocytes were identified by gating on the Hoechst-negative population.

Cells were sorted using a BD Influx Cell Sorter (BD Biosciences). BEL-A CRISPR edited populations were single cell sorted based on viability (DRAQ7 negativity). Primary GLUT1 KO cultures were sorted on days 6 or 7 of differentiation using DRAQ7 and the eGFP-fused GLUT1.RBD to purify the negative population with a gate based on non-targeting guide control cells.

### Osmotic Fragility Assays

Filtered reticulocytes (1-2 ×10^5^/well) were incubated in decreasing NaCl concentrations (0.9–0%) for 10min at 37°C. Lysis was stopped by adding 4x volume of PBSAG. Live cells, considered as having a normal FSC/SSC profile as defined by the 0.9% NaCl control, were counted by flow cytometry using the MACSQuant10^20^.

### Automated Rheoscopy

A total of 1×10^6^ cells were diluted in 200μL of a polyvinylpyrrolidone solution (viscosity, 28.1mPa·s; Mechatronics Instruments). Cell deformability distributions were assessed in an Automated Rheoscope and Cell Analyzer (ARCA) according to previously published protocols^21^. At least 2000 valid cells per sample were analysed.

### Lipid peroxidation

The assay was performed on pre-filtered CD34^+^ cell-derived reticulocytes, in culturing media, using Hoechst to identify reticulocytes. C11-Bodipy (10μM from an DMSO stock, Invitrogen) was used as a fluorescent lipid peroxidation reporter molecule. Oxidation was induced by cumene-hydroperoxide (25μM from an ethanol stock, Sigma). Ethanol was used as a vehicle control. The three reagents were added simultaneously and incubated for 30min at 37°C. Samples were washed 3x with PBS-AG and lipid peroxidation was measured on MACSQuant10 (488 nm).

### Proteomics, Metabolomics, and Lipidomics

Samples were extracted and treated as extensively reported in the Supplementary Methods which also includes details of the Database Searches and Protein Identification following proteomics mass spectrometry. Proteomics^22^ and metabolomics analyses were performed as previously described^23^. Total lipids were extracted as previously described^24^:

## Results

To explore the impact of complete loss of GLUT1 on human erythroid cells, CRISPR mediated gene editing of the BEL-A cell line was first exploited to generate GLUT1 knockdown (KD) and knockout (KO) erythroblast cell lines. Disruptive mono or biallelic mutations respectively were identified in clonal lines (Figure 1A and *Supplemental Figure S1*). GLUT1 expression on expanding cells was measured by flow cytometry using a GLUT1 specific viral receptor binding domain (RBD)^25,26^. Figure 1B confirms a complete loss-of-expression in BEL-A KO cells, while the KD had no overall reduction of GLUT1 when compared with unedited BEL-A (Ctrl). Real-time qPCR (*Supplemental figure S2*) revealed a 70% increase in GLUT1 mRNA on the KD, and a 90% decrease for the KO. RNA levels of an alternative glucose transporter GLUT3, previously reported to be expressed in erythroblasts were significantly upregulated in GLUT1-KO (8-fold upregulation) and KD (2.6-fold increase) respectively.

**Figure 1.**
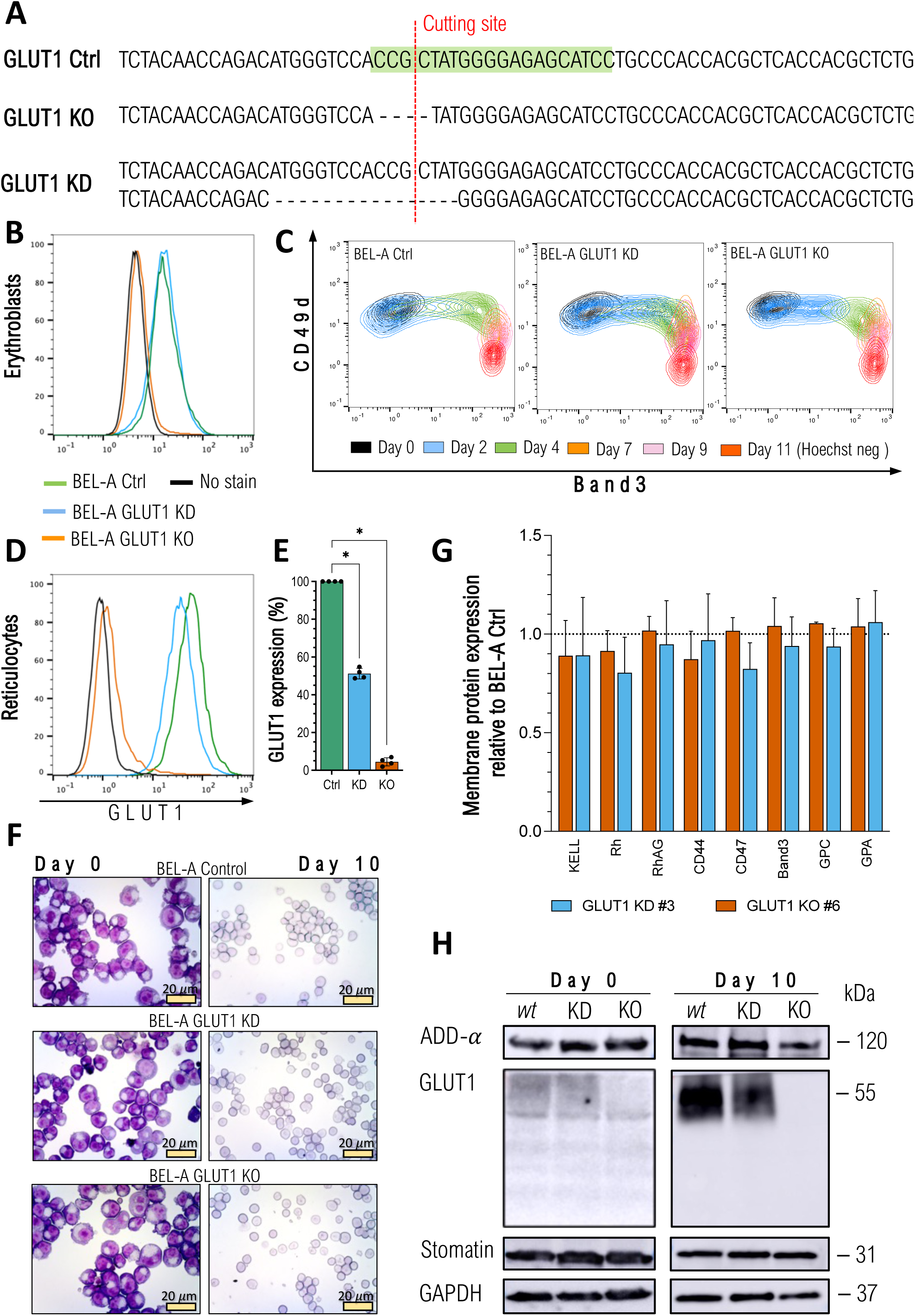
CRISPR-mediated GLUT1 KO on immortalised BEL-A erythroblasts successfully generate reticulocytes. A Sequencing of SLC2A1 (GLUT1) gene on Ctrl BEL-A, highlighting the gRNA (green) used for CRISPR editing. Sanger sequencing of clonal edited lines shows a homozygous 4bp frameshift mutation on the GLUT1 KO line, and a heterozygotic 16bp deletion on the KD line, both in the vicinity of the cutting site (red line). Flow cytometry histogram of GLUT1 staining in BEL-A erythroblasts (B) and derived reticulocytes (D) from ctrl (green), GLUT1 KD (blue), and GLUT1 KO (orange) cell lines compared with no-stain control (black). Cells were stained with anti-GLUT1 FITC conjugate. C Flow cytometry analysis of cell surface marker expression during differentiation. Cells were co-labelled with anti-band3 primary antibody used in conjunction with an IgG1 APC secondary and anti-α4-integrin FITC conjugate. For day 11, reticulocytes were identified using Hoechst as a nuclear DNA stain. E Bar graph illustrates the percentage GLUT1 expression on reticulocytes derived from indicated cell lines. Data is normalised to endogenous expression of Ctrl BEL-A from the median fluorescence intensity (n=4). Individual data points are shown. Error bars indicate standard error of mean. F Representative images of May-Grünwald and Giemsa-stained cytospins depicting expanding BEL-A erythroblasts (Day 0) and corresponding filtered reticulocytes after 10 day differentiation protocol. 40X magnification. Scale bars = 20μm, shown for each image. G Bar graphs illustrate expression of various membrane proteins on reticulocytes derived from indicated cell lines (n=3). Reticulocytes were identified based on Hoechst negativity. Data is normalised to endogenous expression of Ctrl BEL-A and represents the median fluorescence intensity (n=3). Individual data points are shown. Error bars indicate standard error of mean. H Western blots of lysates obtained from indicated cell lines at day 0 of differentiation and reticulocytes filtered after 10 day protocol, incubated with antibodies to α-adducin, GLUT1, Stomatin, and GAPDH (loading control). Multiple Mann-Whitney U tests were used to test for differences between groups. *p < 0.05. Error bars indicate standard deviation.

Since GLUT1 expression increases drastically during erythroid differentiation^27^, the ability of GLUT1-KO erythroblasts to undergo terminal erythroid differentiation and enucleation was assessed through monitoring of Band 3 and α4-integrin (CD49d)^28^. Figure 1C illustrates the ability of both GLUT1 KD and KO lines to differentiate producing the same cascading pattern as the BEL-A Ctrl. GLUT1 expression was assessed on both differentiating erythroblasts and enucleated reticulocytes (Figure 1D-E). Although there was no observed reduction in expression at the erythroblast stage, the heterozygous GLUT1 mutation produced reticulocytes with a 49.2% decrease in surface GLUT1. May-Grünwald and Giemsa stained cytospins of expanding erythroblasts and reticulocytes (Figure 1F) did not identify discernible cellular morphological differences arising from GLUT1 deficiency or absence. Figure 1G illustrates no reductions in surface expression of prominent membrane proteins compared to Ctrl. Complete absence or reduction of GLUT1 was confirmed by immunoblotting and further validated through proteomics (*Supplemental figure S3*). Reported GLUT1 interacting proteins Adducin-α and Stomatin were unaltered in expression (Figure 1H). GLUT4 was detected through proteomics despite its low abundance, with no increased expression in either KD or KO. Conversely, GLUT3 was absent in the proteomic data.

To determine whether compensation for GLUT1 absence is enabled by prolonged expansion of the BEL-A cell line, CRISPR KO was next performed on cultured primary hematopoietic progenitors nucleofected 3 days post isolation as indicated (Figure 2A). Cells were cultured for 21 days and GLUT1 expression assessed by flow cytometry throughout differentiation (Figure 2B). Note that 48h after nucleofection (day 5) it is already possible to discriminate between NT controls and GLUT1-targeted populations with clear negative comprising approximately 75% of the total cells discernible on day 6.

**Figure 2.**
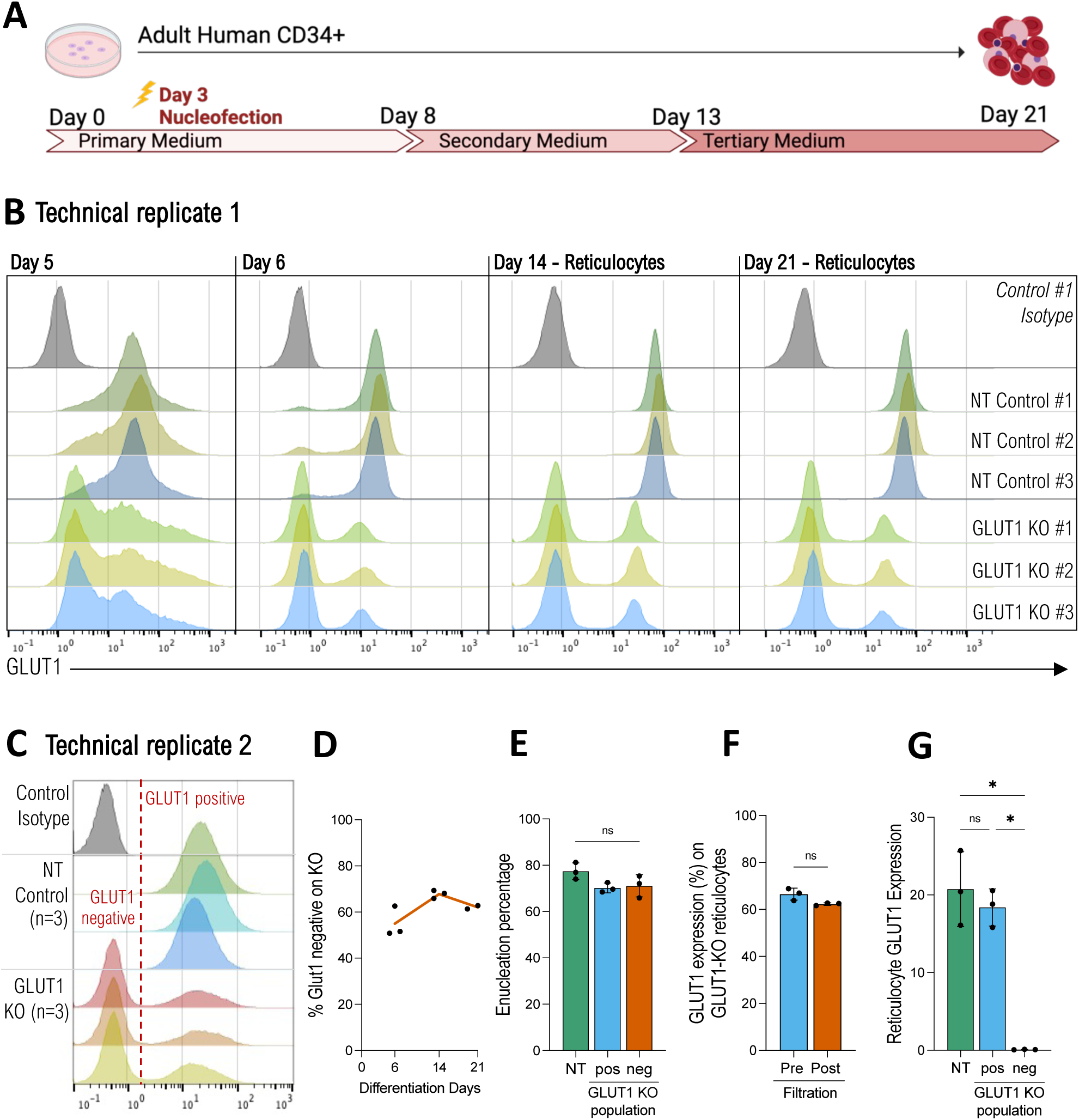
Primary erythropoiesis is not affected by knocking out GLUT1. A Schematic diagram of human CD34+ 3-step culture method. PBMCs are isolated from apheresis cones, followed by CD34+ magnetic separation. Cells are then nucleofected on day 3 with Non-Targeting (NT) or GLUT1 specific gRNAs. B Flow cytometry histograms show GLUT1 expression of 3 donors, nucleofected with NT or SLC2A1 targeting gRNAs, on days 5, 6, 14, and 21 of differentiation. For days 14 and 21 Hoechst was used to identify the reticulocytes. Cells were stained with anti-GLUT1 FITC conjugate (n=3) or a no-stain control (black). C Day 21 filtered reticulocytes stained with anti-GLUT1 FITC conjugate (n=3) or a no-stain control (black). D Percentage of GLUT1 negative population on GLUT1-targeted KO (n=3) on days 6, 14 and 21 of differentiation. E Percentage of reticulocytes (Hoechst negative) at day 21 of differentiation on NT control and negative and positive GLUT1 populations of the GLUT1-targeted KO. F Percentage of GLUT1 negative population on GLUT1-targeted KO on reticulocytes pre- and post-filtration. G Bar graph illustrates GLUT1 expression on reticulocytes from NT and GLUT1 negative and positive populations of the GLUT1-targeted KO. Data represents the median fluorescence intensity (n=3). Individual data points are shown. Multiple Mann-Whitney U tests were used to test for differences between groups. *p < 0.05. Error bars indicate standard deviation.

GLUT1 expression was measured on Hoechst negative reticulocytes present at days 14 and 21 with the same proportion (75%) of cells shown to be GLUT1 negative as on day 6. Of note, assessment of the GLUT1 positive population within the CRISPR-edited samples reveals a decreased GLUT1 expression when compared with the matched NT control, signifying a knockdown population. It is noteworthy that cells from all three donors enucleated and from day 6 of erythroid differentiation maintained a stable GLUT1-negative population, indicating no competitive disadvantage in erythroblast expansion resulting from absence of GLUT1.

We performed a replicate assay of the CRISPR-KO (Figure 2C-G), resulting in an average of 62% GLUT1-negative population across 3 donors. Here, within the GLUT1 positive population, expression of GLUT1 is comparable to the matched NT controls. By co-labelling day 21 pre-filtration samples with anti-GLUT1.RBD and Hoechst it was possible to compare enucleation of GLUT1-positive and -negative populations within GLUT1-targeted cultures, as well as with the NT controls (Figure 2E). There is no significant difference between the 3 populations demonstrating that the absence of GLUT1 does not adversely affect the membrane remodelling and cytoskeletal changes required for nucleus extrusion. To assess the effect of GLUT1 absence on physical membrane properties of reticulocytes day 21 cultures were filtered using a 5μm filter with the GLUT1-negative cells of each KO-targeted donor assessed before and after filtration to establish differential ability to pass through the filter (Figure 2F). No significant difference in proportion of GLUT1-negative cells pre- and post-filtration was observed (66.5+/-2.6% vs 62.3+/-0.6%) indicating that absence of GLUT1 does not impact reticulocyte capacity to traverse 5μm pores.

To further investigate the properties of GLUT1-negative reticulocytes, additional knockouts were performed and enriched for using FACS. As previously illustrated for the BEL-A GLUT1 KO, no substantial differences were identified in terminal erythroid differentiation per CD49d/Band 3 labelling (Figure 3A), nor did cell morphology change between CD34+ NT and GLUT1-KO populations (Figure 3B). Expression of surface membrane proteins was analysed by flow cytometry, with the median fluorescence intensity averaged for the NT controls (Figure 3C). Complete absence of GLUT1 surface expression was confirmed, and no alterations in expression of most major erythrocyte membrane proteins as indicated was observed. Within the NT controls, the expression of BCAM (Basal Cell Adhesion Molecule, or Lutheran blood group protein^29^) exhibits considerable variability due to a well-established donor variable dual population of Lu presenting cells (*Supplemental figure S4*). However, even considering this variability, there remains a statistically significant reduction in BCAM expression in the GLUT1-KO compared with the matched NT cultures. Also, significantly reduced, albeit mildly, are CD47, Rh, and RhAG (components of the Rh subcomplex).

**Figure 3.**
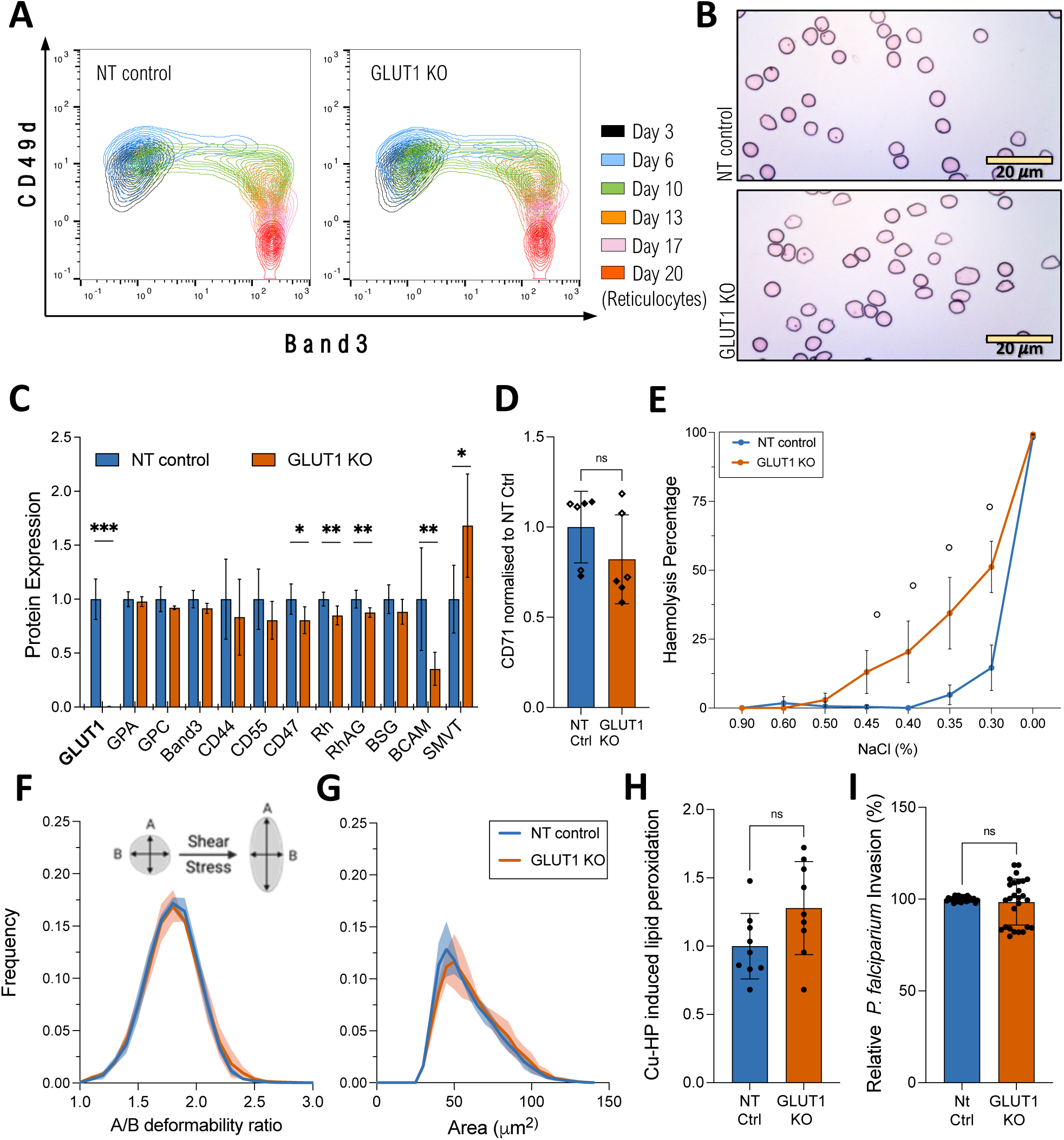
Properties of CD34+ GLUT1-KO derived reticulocytes. A Waterfall indicating progression of control and GLUT1-KO transfected cultures of single donor CD34+ differentiation. Cells were co-labelled with anti-band3 primary antibody used in conjunction with an IgG1 APC secondary and anti-α4-integrin FITC conjugate. For day 20, reticulocytes were identified using Hoechst as a nuclear DNA stain. B Representative cytospins of the same culture were obtained on day 20 post-leukofiltration. 40X magnification. Scale bars = 20μm, shown for each image. C Bar graphs indicate the expression of various membrane proteins on reticulocytes derived from NT or GLUT1-KO primary cultures (n=6, for SMVT n=3, with 2 technical repeats each). D Anti-transferrin receptor (CD71) labelling of on reticulocytes (n=6, open or filled dots indicate 2 separate cultures, each with 3 donors). For both, reticulocytes were leukofiltered on day 20. Data is normalised to endogenous expression of NT control and represents the median fluorescence intensity. E Osmotic resistance analysis calculated based on viable cell counts (flow cytometer) after incubation with decreasing concentrations of NaCl (n=3, 2 technical replicates, ∘ p≤0.0021). F Deformability and G cell area (μm^2^) were measured under shear stress by Automated Rheoscope Cell Analyser (n=3, N>2000 cells), which elongates cells and measures length over width as deformability parameter. Shaded region represents the standard deviation. H Quantitative analysis of lipid peroxidation detected by a shift in the fluorescence signal after treatment with 25mM cumene hydroperoxide. Data normalized to NT-control (n=3, 3 technical replicates) I Bar graph quantifying *Plasmodium falciparum* reticulocyte invasion efficiency. Invasion was assessed by flow cytometry using a SYBR-green DNA stain (3 separate parasitaemia percentages, 3 technical replicates) and data was normalised *per* donor (n=3) to the NT-control. Figure 3 comprises data obtained from two independent cultures, each of 3 donors with figures 3A and B presenting representative data from cultures with FACS sorted GLUT1-KO purity of 99%, E-I 94% and C-D combined data from all 6 donors. A non-parametric Mann-Whitney U test or Kruskal-Wallis test with Bonferroni correction were used to test for differences between groups. *p < 0.05, **p < 0.01, ***p < 0.001. Error bars indicate standard deviation.

Expression of various cell surface nutrient transporters was also measured, using RBDs that function as specific ligands of solute carrier (SLC) nutrient transporters^30^. Amongst these, the sodium dependent multivitamin transporter (SMVT; product of the SLC5A6 gene) is the only protein significantly increased in the GLUT1-KO samples using available reagents. Data on the remaining nutrient transporters can be seen in *Supplemental figure S5*. Reticulocyte maturity as assessed by CD71^31^ (transferrin receptor) levels varied between cultures but was not consistently or significantly altered by the absence of GLUT1(Figure 3D).

To provide a global overview of alterations to protein expression, matched NT-control and GLUT1-KO reticulocytes (n=3) were subjected to proteomic analysis (Figure 4). Deficiency of GLUT1 was confirmed in GLUT1-KO, with a 26-fold change decrease in the GLUT1-KO reticulocytes compared to NT-controls (Figure 4B, p < 0.001). GLUT4 was detected despite its low abundance but showed no increased expression in the KOs. No specific vitamin C or DHA transporters were identified, nor was SMVT detected in the data set. Proteins known to interact with GLUT1, α- and β-adducin, dematin, and stomatin, show no significant alteration in expression in the KO samples. CD47, Rh (RhD and RhCE), RhAG, and BCAM also show no significant changes in total protein content despite the reduction seen in surface levels. However, gene ontology enrichment analysis revealed a highly significant enrichment of glycolysis-associated proteins (P<10-10), with downregulation of the entire glycolytic pathway, and conversely an upregulation of AMPK-signalling and autophagy-associated proteins, promoting catabolic pathways to generate more ATP.

**Figure 4.**
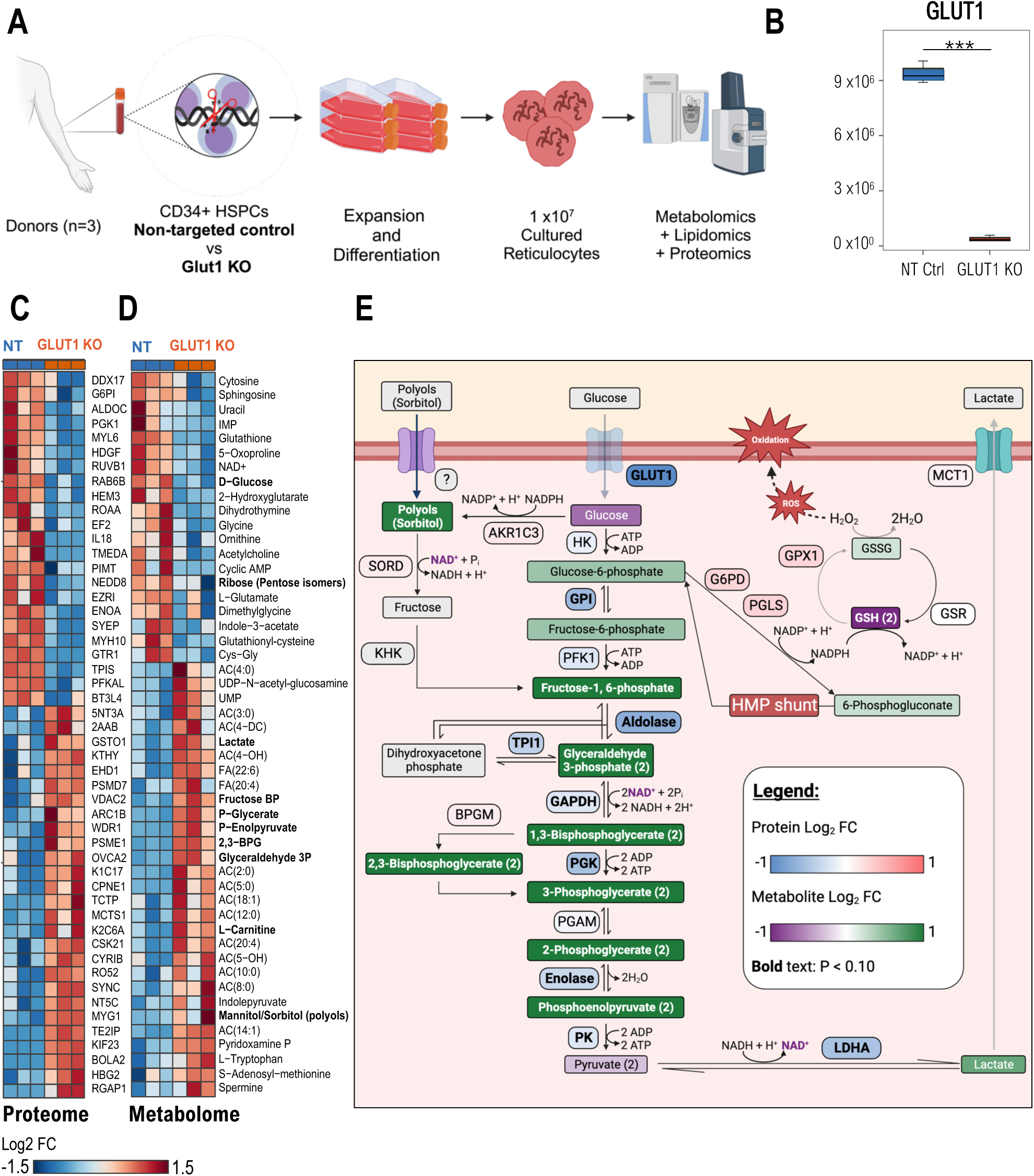
Proteomics confirms GLUT1 absence in CD34+ GLUT1-KO derived reticulocytes and Metabolite analyses reveals downregulated metabolic processes. A Simplified schematic of multi-omics cell preparation, where CD34+ from 3 donors were nucleofected with either GLUT1-or non-targeting gRNAs, expanded and differentiated into reticulocytes. 10 million filtered reticulocytes were needed for comprehensive analyses of the proteome, metabolome, and lipidome. Made using BioRender. B Box plot comparing GLUT1 protein level between NT and GLUT1 KO reticulocytes as Log_2_(Fold change). Box plot analysis (mean±min to max with standard deviation) was performed by RStudio, and significance was calculated upon false discovery rate correction (***p < 0.001).C,D Hierarchical clustering of the Top 50 t-test significant proteins (C) and metabolites (D) between NT and GLUT1 KO CD34^+^-derived reticulocytes. E Schematic representation of Glycolysis, the Polyol pathway, and the Glutathione redox cycle where proteins and metabolites are colour-coded by Log_2_ (Fold Change) of GLUT1-KO reticulocytes in relation to NT control. The 10 steps of glycolysis are represented, with glucose and lactate both reduced in the KO while the remaining intermediate products increased. All involved enzymes are decreased (HK, Hexokinase; GPI, Glucose-6-phosphate isomerase; PFK1, Phosphofructokinase-1; TPI1, Triosephosphate isomerase; GAPDH, Glyceraldehyde-3-phosphate dehydrogenase; BPGM, Biphosphoglycerate mutase; PGK, Phosphoglycerate kinase; PGAM, Phosphoglycerate mutase; PK, Pyruvate kinase; LDHA, Lactate dehydrogenase A). There is an imbalance in the Glutathione cycle, as a consequence of increased ROS, characterised by the depletion of the reduced form (GSH) and increase of oxidised glutathione (GSSG) and glutathione peroxidase 1 (GPX1). The Hexose monophosphate (HMP) shunt is upregulated as source of NADPH, needed to maintain glutathione in its reduced form. (G6PD, Glucose-6-phosphate dehydrogenase; PGLS, 6-phosphogluconolactonase; GSR, Glutathione-disulfide reductase). An increase in polyols (such as sorbitol and mannitol) was also detected, which can be converted into Fructose-1,6-phosphate by sorbitol dehydrogenase (SORD) and ketohexokinase (KHK). Created with Biorender.

*In vivo*, RBCs remain in circulation for up to 120 days, being continuously exposed to shear-stress and repeated elastic deformations. Given the varied potential contributions to RBC stability that arise from GLUT1 influence on membrane-cytoskeletal connectivity, solute transport, and metabolism we employed a variety of *in vitro* techniques designed to dissect the effects of absence of GLUT1 in reticulocytes. Osmotic fragility assays summarised in Figure 3E illustrate increased haemolysis compared to control cells evident at 0.45% NaCl and maintained at lower NaCl levels, indicative of a higher osmotic fragility on GLUT1-KO cells and suggestive of a potential defect in volume homeostasis or ion balance.

GLUT1, via association with adducin and dematin is reported to provide a site of vertical attachment between membrane and cytoskeleton in red blood cells. To investigate cell deformability and membrane stability cells were subjected to Automated Rheoscopy (ARCA^32^. Figures 3F-G indicate no differences in cross sectional area or capacity to undergo deformation (elongation index) arising from the absence of GLUT1 illustrating no detectable reductions in deformability.

In addition to its role in glucose transport, GLUT1 also acts as a transporter of DHA which gets converted into ascorbic acid in the cytoplasm, a potent antioxidant^33^. Using C11-Bodipy as a lipid peroxidation sensor we measured ROS response using cumene hydroperoxide (Cu-HP) as an oxidising agent. Although Figure 3H shows a small increase in lipid peroxidation (p=0.0726) in the KO population, the range of individual data points precludes it from reaching statistical significance. As an additional assay to exploit the generation of this novel GLUT1 KO cell resource reticulocytes were subjected to invasion assays with the malaria parasite *Plasmodium falciparum*. Interestingly, despite the abundance of GLUT1 within the reticulocyte/erythrocyte membrane, absence of GLUT1 did not impair the ability of reticulocytes to support invasion by *P. falciparum* with no significant differences in invasion efficiency of GLUT1 KO compared to NT control reticulocytes (Figure 3I), indicative that GLUT1 is not exploited as an essential receptor for merozoite invasion.

The apparent absence of phenotype arising from the loss of such an upregulated and abundantly expressed protein within the erythroid lineage was surprising. To further explore potential distinctions that arise from the absence of GLUT1, the samples already tested for proteomic differences were also investigated to uncover metabolomic and lipidomic disparities (Figures 4 and 5). Interestingly, metabolomics analyses revealed that GLUT1-KO reticulocytes do compensate for a decrease in glucose uptake, as inferred by a significantly lower steady state levels of glucose and by up-regulating glucose catabolism downstream to hexose phosphate. Figure 4D summarises the metabolomic and proteomic alterations observed in the GLUT1-KO. Indeed, in GLUT1-KO cells we detected significant elevation in the levels of fructose bisphosphate, glyceraldehyde 3-phosphate, 2,3-bisphosphoglycerate (BPG), phosphoglycerate (2 and 3 isomers), and phosphoenolpyruvate. However, despite net significant increases in lactate, decreases in pyruvate were observed, suggestive of altered pyruvate to lactate ratios, perhaps because of increased NADH-dependent methaemoglobin reductase activity, which could outcompete lactate dehydrogenase for the same cofactor. Despite higher levels of virtually all glycolytic intermediates, especially at the payoff steps of glycolysis, no significant changes were observed between the two groups, suggesting that, in the absence of perturbation, increased glycolytic fluxes may compensate for decrease glucose uptake in the KO reticulocytes. Interestingly, elevation in polyols such as sorbitol which can enter glycolysis at the fructose bisphosphate level are suggestive of alternative sugar substrates that could contribute to compensating for the partially ablated glucose uptake.

**Figure 5.**
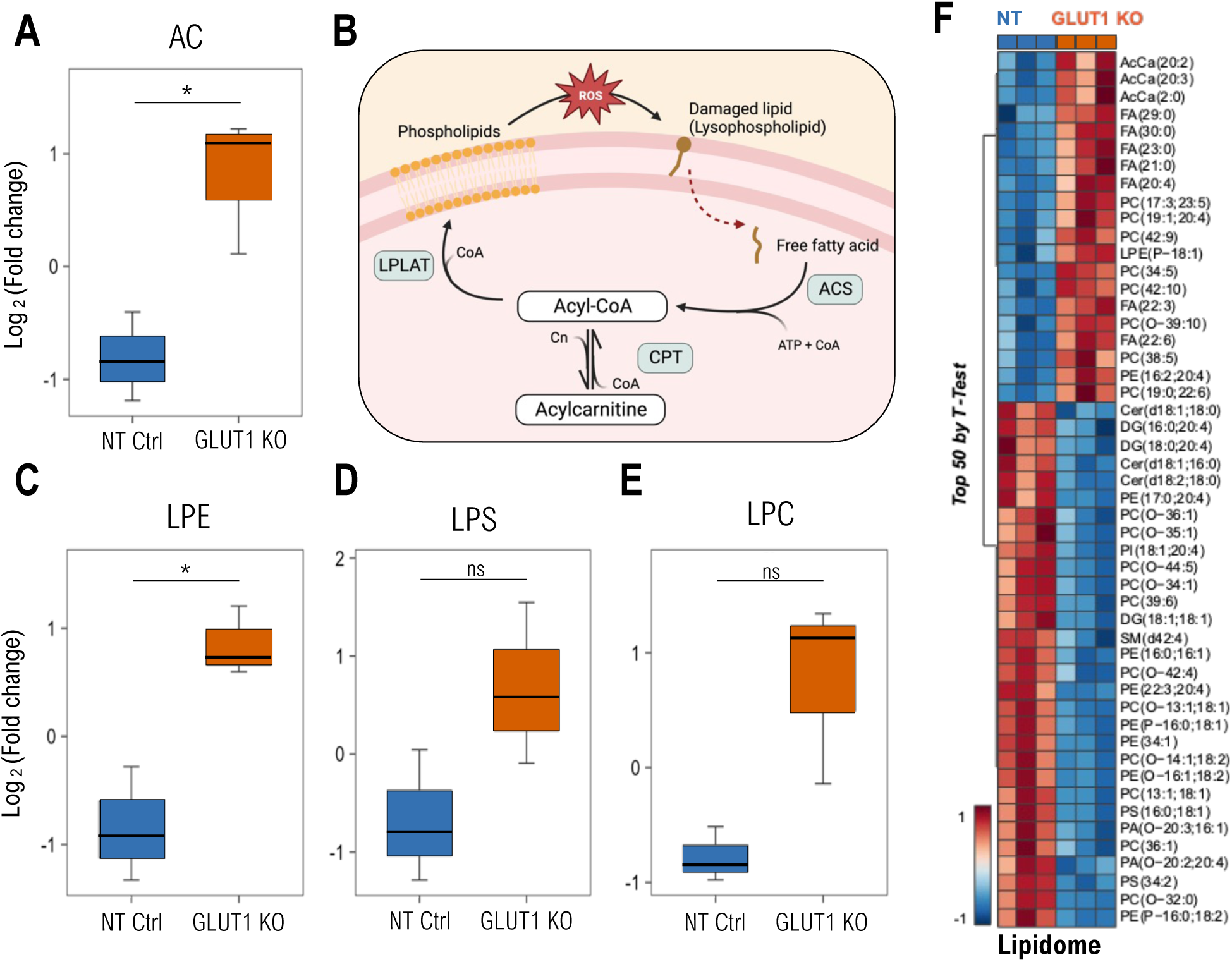
Lipid composition highlights increased oxidant stress to the membrane of GLUT1-KO reticulocytes. A Box plot showing the comparison of Acyl-carnitines as Log_2_(FoldChange, FC), between NT and GLUT1-KO reticulocytes. Box plots analysis (mean±min to max with standard deviation) was performed by RStudio, and significance was calculated upon false discovery rate correction (p < 0.05). B Simplified schematics of the Lands Cycle, capable of repairing damaged lipids (lysophospholipids) generated from increased oxidative stress. The damaged chain is removed by a phospholipase, originating a free fatty acid which is converted into Acyl-CoAs by Acyl-CoA synthetase (ACS) in an ATP-dependent reaction. Acyl-CoAs can be converted into Acyl-carnitines by carnitine palmitoyltransferase (CPT). Lysophospholipid acyltransferases (LPLATs) incorporate undamaged fatty acid chains into membrane lysophospholipids, repairing the lipid membrane. Created in Biorender. C-E Box plots comparing Lysophosphatidylethanolamine (LPE, C), Lysophosphatidylserines (LPS, D) and Lysophosphatidylcholine (LPC, E) between NT and GLUT1-KO reticulocytes, presented as Log^2^(FC). F Hierarchical clustering of the Top 50 t-test significant lipids between NT and GLUT1 KO CD34^+^-derived reticulocytes.

Despite these adaptations, markers of oxidant stress were observed in the GLUT1-KO group, even in the absence of perturbation. For example, KO cells showed significantly lower levels of reduced glutathione (GSH), with near significant (p = 0.06) elevation in oxidized to reduced glutathione ratios (*Supplemental Figure S6*), suggestive of a rewiring of glucose consumption mainly through glycolysis, perhaps via the reduction of glucose oxidation via the pentose phosphate, a key pathway for the generation of the reducing equivalent NADPH, which is directly or indirectly involved in virtually all antioxidant reactions in the mature erythrocyte.

Finally, and most evidently, L-carnitine and all acyl-carnitines (AC) were found to increase in the KO reticulocytes (Figure 5A). In mature RBCs this class of metabolites is associated with lipid membrane damage repair via the Lands cycle^34^, represented in Figure 5B. Lipidomics results indicated significant decreases in the levels of cholesteryl-esters (ChE), diacylglycerols (DG), ceramides (Cer) and all phospholipids (PC, PE, PI, PS, PG, PA) in the KO group, except for triacylglycerols (TG), free fatty acids (FFA), and mono-hexosylceramides (Hex1Cer) – *Supplemental figure S7*. Most importantly, elevation in all lysophospholipids (significant for LPE, borderline significant for LPS and LPC, as seen in Figures 5C-E) are suggestive of increased phospholipase activity upon activation of the Lands cycle for the repair of damaged lipids in the KO group.

## Discussion

As one of the most abundantly expressed proteins within the erythrocyte membrane, GLUT1 is widely assumed to play a crucial role in the development, structure, and function of the red blood cell. However, suitable erythroid models to study the effect of lack of GLUT1 in humans is lacking. Mouse models are inappropriate owing to the absence of GLUT1 in the adult mouse erythrocyte^40^ and to date in human erythroid cells only a 60% knockdown has been achieved^41^, levels that mimic the naturally occurring GLUT1 deficiency syndrome^26^, a disease that is haematologically symptomless despite presentation of neurological phenotypes. Here, we demonstrate the ability to generate completely GLUT1 deficient erythroid cells that successfully complete terminal differentiation to generate enucleated reticulocytes unexpectedly deficient in phenotype, rewriting our understanding of the essentiality of this protein in human erythroid biology. RBC structural integrity and capacity for deformation arise from vertical protein associations between integral membrane proteins and the underlying spectrin-based cytoskeleton, mediated via an array of cytoskeletal adaptor proteins. It would not be unreasonable to hypothesise that loss of GLUT1, a protein that accounts for around 10% of the protein component of the RBC membrane^42^ would confer disruption to membrane integrity. Surprisingly, however, there were no reductions in reported GLUT1 associating proteins, nor any other detectable membrane or cytoskeletal protein were observed (aside from a mild reduction in Rh subcomplex components). Furthermore, neither deformability, as assessed through rheoscopy, nor ability to support invasion by *P. falciparum* was found to be altered in GLUT1 KO reticulocytes. No significant compensatory upregulation of membrane proteins that could be anticipated to occupy vacant binding sites or residency within the plasma membrane was observed through proteomics, indicating that GLUT1 does not play a direct role in maintenance of the structural integrity of the RBC via its protein-protein interactions.

Metabolomic studies revealed that GLUT1-KO reticulocytes exhibit hallmarks indicative of reduced glucose import with downregulated metabolic processes and upregulated AMPK signalling, consistent with reduced glucose content. Surprisingly, the absence of GLUT1 does not present an obvious impediment to erythroblast terminal differentiation or enucleation; this may reflect initial compensation of glucose import mediated via GLUT3, which has been previously demonstrated to be expressed in the early stages of differentiation before loss and replacement by GLUT1^27^. However, there was no upregulation of alternative glucose transporters such as GLUT3 or GLUT4 in differentiated reticulocytes, nor increased abundance of mitochondria to maximise energy output, as inferred from comparable levels of mitochondrial proteins via proteomics between NT and GLUT1-KO cells. Thus, our data show for the first time that GLUT1 is not required for erythroid cells to obtain sufficient glucose to drive terminal differentiation.

Whilst GLUT1 is amongst the proteins most upregulated during erythropoiesis, seminal work from Montel-Hagen et al^9^ demonstrated that glucose transport actually decreases during the latter stages of erythroid differentiation, replaced instead by transport of L-dehydroascorbic acid (DHA), an oxidised form of ascorbic acid in a switch regulated by stomatin association. Absence of GLUT1 is therefore predicted to also impact intracellular antioxidant levels and redox balance. Interestingly, contrastingly to the unaltered deformability, increased osmotic fragility of GLUT1-KO reticulocytes compared to unedited controls was detected, mimicking a phenotype observed in RBCs of mice with reduced ascorbic acid intake ^37,43^. Metabolomics also revealed a depletion of the glutathione pool. These changes were accompanied by the maintenance of total ATP levels with increased steady state levels of glycolysis intermediates downstream to fructose bisphosphate, elevation in the levels of alternative sugar substrates like sorbitol and decrease in the levels of pentose phosphate pathway metabolites.

Absence of GLUT1 mediated DHA transport likely adds pressure to the remaining cellular antioxidant defence mechanisms while trying to maintain redox balance, as suggested by the borderline (p = 0.07) decreases in DHA in the KO cells. Of note, in a background of minimal detectable compensatory alterations we did observe increased expression of the sodium dependent multivitamin transporter (SMVT) which catalyses the uptake of α-lipoic acid, a potent antioxidant, pantothenic acid (vitamin B5 – precursor to CoA that participates in heme synthesis^44^ and the Lands cycle), and biotin^45^ in primary GLUT1-KO reticulocytes. Our data indicate a mild and potentially partially compensated defect in antioxidant capacity of GLUT1 KO reticulocytes.

Altogether, these results indicate that, while non-lethal in the absence of perturbations, the GLUT1 KO mutation introduces a strain to RBC antioxidant metabolism by forcing glycolytic metabolism through the Embden-Meyerhof Parnas pathway at the expense of antioxidant pathways such as the hexose monophosphate shunt. Interestingly, these metabolic alterations were accompanied by elevation in acyl-carnitine pools, lysophospholipids, and odd-chain fatty acids derived from alpha-oxidation of longer chain polyunsaturated fatty acids, despite decreases in several classes of lipids. These lipidomics data are suggestive of increased oxidant stress to the RBC membrane fraction, consistent with the activation of the Lands cycle and an increased susceptibility to osmotic fragility, similar to that observed for RBCs from patients suffering from chronic^46^ or acute kidney disease^47,48^, sickle cell disease and hypoxia^49^.

*In vivo* circulating RBCs are highly exposed to oxidative stress with repetitive nature of erythrocyte function hard to replicate *in vitro*, such as the accumulation of reactive oxygen species and consequent repetitive deformability decreases. Whilst experimentally impractical to explore, we recognise that as reticulocytes derived through *in vitro* culture, the full effects of complete GLUT1 deficiency subject to the rigours of *in vivo* circulation and in the absence of residual mitochondria (loss of which is induced by circulation) may not be evident. We predict that GLUT1-KO will most likely exhibit a reduced *in vivo* circulatory half-life as a consequence of increased osmotic fragility, altered energy metabolism, and disrupted redox balance observed here.

These data substantially extend our understanding of the role glucose transport and metabolism play in erythropoiesis and redefines the importance of GLUT1 in RBC membrane structure. Furthermore, for many years, the absence of haematological phenotype associated with reduced GLUT1 expression in G1DS has remained unexplained, attributed to the remaining copies of the transporter still present. In generating completely deficient GLUT1 erythroid cells that demonstrate no apparent defects in expansion, enucleation, or structural phenotypes, we now provide cell biological evidence that supports the clinical consensus that a reduction in GLUT1 abundance in RBCs does not cause anaemia in G1DS.

## Author contributions

CMF, AMT and TJS conceived and designed the study. CMF performed and analysed the majority of experiments, NRK and TJS contributed to experimental work and analysis. JGGD and GJS developed ARCA hardware and analysis software, PM performed proteomics analysis, MD, DS, AD, and PM conducted omics analyses and interpretation, AMT and TJS supervised the study. CMF, AMT and TJS wrote the manuscript. All authors reviewed and approved the final manuscript.

## Acknowledgements

The authors wish to thank Andrew Herman from the University of Bristol Faculty of Biomedical Sciences Flow Cytometry Facility for cell sorting support and Vincent Petit (Metafora Biosystems) for kind provision of anti-GLUT1 and remaining SLC-targeting RBDs. This study was supported through funding provided by the European Union ITN ‘EVIDENCE’ grant agreement ID 860436 for CMF, the Medical Research Council (MR/V010506/1) for TJS and infrastructure support funding from the National Institute for Health Research Blood and Transplant Research Unit (NIHR BTRU) in Red Cell Products (IS-BTU-1214-10032). PLM is supported by grants from the Swedish Cancer Foundation / Cancerfonden (21 0340) and from the Myelodysplastic Syndromes Foundation, Inc. (1142079). AD was supported by funds from the National Heart, Lung and Blood Institutes (NHLBI) R01 HL146442, R01 HL161004, R01 HL148151, R21 HL150032

The views expressed are those of the authors and not necessarily those of the National Health Service, NIHR, or the Department of Health and Social Care

## References

1. Mueckler M, Caruso C, Baldwin SA, et al. Sequence and structure of a human glucose transporter. Science. 1985;229(4717):941–945.

2. Montel-Hagen A, Blanc L, Boyer-Clavel M, et al. The Glut1 and Glut4 glucose transporters are differentially expressed during perinatal and postnatal erythropoiesis. Blood. 2008;112(12):4729–4738.

3. Li H, Lykotrafitis G. Erythrocyte membrane model with explicit description of the lipid bilayer and the spectrin network. Biophys J. 2014;107(3):642–653.

4. Mohandas N, Gallagher PG. Red cell membrane: past, present, and future. Blood. 2008;112(10):3939–3948.

5. Guizouarn H, Allegrini B. Erythroid glucose transport in health and disease. Pflugers Arch. 2020;472(9):1371–1383.

6. Van Wijk R, Van Solinge WW. The energy-less red blood cell is lost: Erythrocyte enzyme abnormalities of glycolysis. Blood. 2005;106(13):4034–4042.

7. Cilek N, Ugurel E, Goksel E, Yalcin O. Signaling mechanisms in red blood cells: A view through the protein phosphorylation and deformability. J Cell Physiol. 2023;

8. Moura PL, Iragorri MAL, Français O, et al. Reticulocyte and red blood cell deformation triggers specific phosphorylation events. Blood Adv. 2019;3(17):2653–2663.

9. Lie Montel-Hagen A, Kinet S, Manel N, et al. Erythrocyte Glut1 Triggers Dehydroascorbic Acid Uptake in Mammals Unable to Synthesize Vitamin C.

10. Yang H, Wang D, Engelstad K, et al. Glut1 deficiency syndrome and erythrocyte glucose uptake assay. Ann Neurol. 2011;70(6):996–1005.

11. Winczewska-Wiktor A, Hoffman-Zacharska D, Starczewska M, et al. Variety of symptoms of GLUT1 deficiency syndrome in three-generation family. Epilepsy & Behavior. 2020;106:107036.

12. Klepper J, Akman C, Armeno M, et al. Glut1 Deficiency Syndrome (Glut1DS): State of the art in 2020 and recommendations of the international Glut1DS study group. Epilepsia Open. 2020;5(3):354–365.

13. Olivotto S, Duse A, Bova SM, et al. Glut1 deficiency syndrome throughout life: clinical phenotypes, intelligence, life achievements and quality of life in familial cases. Orphanet J Rare Dis. 2022;17(1):1–13.

14. Furuse T, Mizuma H, Hirose Y, et al. A new mouse model of GLUT1 deficiency syndrome exhibits abnormal sleep-wake patterns and alterations of glucose kinetics in the brain. DMM Disease Models and Mechanisms. 2019;12(9):.

15. Willemsen MA. Anemia in Glucose Transporter Type 1 Deficiency Syndrome: Often Expected, Rarely Encountered, and with a Fascinating Explanation. Neuropediatrics. 2017;48(5):327–328.

16. Satchwell TJ, Wright KE, Haydn-Smith KL, et al. Genetic manipulation of cell line derived reticulocytes enables dissection of host malaria invasion requirements. Nat Commun. 2019;10(1):.

17. Trakarnsanga K, Griffiths RE, Wilson MC, et al. An immortalized adult human erythroid line facilitates sustainable and scalable generation of functional red cells. Nat Commun. 2017;8(1):1–7.

18. Hawksworth J, Satchwell TJ, Meinders M, et al. Enhancement of red blood cell transfusion compatibility using CRISPR-mediated erythroblast gene editing. EMBO Mol Med. 2018;10(6):e8454.

19. King NR, Martins Freire C, Touhami J, et al. Basigin mediation of Plasmodium falciparum red blood cell invasion does not require its interaction with monocarboxylate transporter 1.

20. Bernecker C, Köfeler H, Pabst G, et al. Cholesterol Deficiency Causes Impaired Osmotic Stability of Cultured Red Blood Cells. Front Physiol. 2019;10:1529.

21. Moura PL, Hawley BR, Mankelow TJ, et al. Non-muscle myosin II drives vesicle loss during human reticulocyte maturation. Haematologica. 2018;103(12):1997–2007.

22. Thomas T, Stefanoni D, Dzieciatkowska M, et al. Evidence for structural protein damage and membrane lipid remodeling in red blood cells from COVID-19 patients. J Proteome Res. 2020;19(11):4455.

23. Nemkov T, Reisz JA, Gehrke S, Hansen KC, D’Alessandro A. High-throughput metabolomics: Isocratic and gradient mass spectrometry-based methods. Methods in Molecular Biology. 2019;1978:13–26.

24. Reisz JA, Zheng C, D’Alessandro A, Nemkov T. Untargeted and semi-targeted lipid analysis of biological samples using mass spectrometry-based metabolomics. Methods in Molecular Biology. 2019;1978:121–135.

25. Kinet S, Swainson L, Lavanya M, et al. Isolated receptor binding domains of HTLV-1 and HTLV-2 envelopes bind Glut-1 on activated CD4+ and CD8+ T cells. Retrovirology. 2007;4:31.

26. Gras D, Cousin C, Kappeler C, et al. A simple blood test expedites the diagnosis of glucose transporter type 1 deficiency syndrome. Ann Neurol. 2017;82(1):133–138.

27. Gautier E-F, Ducamp S, Leduc M, Zermati Y, Correspondence PM. Comprehensive Proteomic Analysis of Human Erythropoiesis. 2016;

28. Hu J, Liu J, Xue F, et al. Isolation and functional characterization of human erythroblasts at distinct stages: implications for understanding of normal and disordered erythropoiesis in vivo. Blood. 2013;121(16):3246–3253.

29. Parsons SF, Lee G, Spring FA, et al. Lutheran blood group glycoprotein and its newly characterized mouse homologue specifically bind α5 chain-containing human laminin with high affinity. Blood. 2001;97(1):312–320.

30. Justus D, Yan H, Hale J, et al. Evaluating Metabolic Regulation of Human Erythropoiesis By Profiling Nutrient Transporter Expression.

31. Malleret B, Xu F, Mohandas N, et al. Significant Biochemical, Biophysical and Metabolic Diversity in Circulating Human Cord Blood Reticulocytes. PLoS One. 2013;8(10):e76062.

32. Moura PL. Investigating the process of reticulocyte maturation to erythrocytes in vitro. 2020;

33. Pallotta V, Gevi F, D’Alessandro A, Zolla L. Storing red blood cells with vitamin C and N-acetylcysteine prevents oxidative stress-related lesions: A metabolomics overview. Blood Transfusion. 2014;12(3):376–387.

34. Nemkov T, Skinner S, Diaw M, et al. Plasma Levels of Acyl-Carnitines and Carboxylic Acids Correlate With Cardiovascular and Kidney Function in Subjects With Sickle Cell Trait. Front Physiol. 2022;13:.

35. D’Alessandro A, Anastasiadi AT, Tzounakas VL, et al. Red Blood Cell Metabolism In Vivo and In Vitro. Metabolites. 2023;13(7):.

36. Gwozdzinski K, Pieniazek A, Gwozdzinski L. Reactive Oxygen Species and Their Involvement in Red Blood Cell Damage in Chronic Kidney Disease. Oxid Med Cell Longev. 2021;2021:.

37. Tu H, Wang Y, Li H, Brinster LR, Levine M. Chemical Transport Knockout for Oxidized Vitamin C, Dehydroascorbic Acid, Reveals Its Functions in vivo. EBioMedicine. 2017;23:125–135.

38. Kodippili GC, Putt KS, Low PS. Evidence for three populations of the glucose transporter in the human erythrocyte membrane. Blood Cells Mol Dis. 2019;77:61–66.

39. Burton NM, Bruce LJ. Modelling the structure of the red cell membrane. Biochemistry and Cell Biology. 2011;89(2):200–215.

40. Montel-Hagen A, Blanc L, Boyer-Clavel M, et al. The Glut1 and Glut4 glucose transporters are differentially expressed during perinatal and postnatal erythropoiesis. Blood. 2008;112(12):4729–4738.

41. Oburoglu L, Tardito S, Fritz V, et al. Glucose and Glutamine Metabolism Regulate Human Hematopoietic Stem Cell Lineage Specification. Cell Stem Cell. 2014;15(2):169–184.

42. Mueckler M. Facilitative glucose transporters. Eur J Biochem. 1994;219(3):713–725.

43. Tu H, Li H, Wang Y, et al. Low Red Blood Cell Vitamin C Concentrations Induce Red Blood Cell Fragility: A Link to Diabetes Via Glucose, Glucose Transporters, and Dehydroascorbic Acid. EBioMedicine. 2015;2(11):1735–1750.

44. Mian SA, Philippe C, Maniati E, et al. Vitamin B5 and succinyl-CoA improve ineffective erythropoiesis in SF3B1 mutated myelodysplasia. Sci Transl Med. 2023;15(685):eabn5135.

45. Quick M, Shi L. The Sodium/Multivitamin Transporter: A Multipotent System with Therapeutic Implications. Vitam Horm. 2015;98:63–100.

46. Xu P, Chen C, Zhang Y, et al. Erythrocyte transglutaminase-2 combats hypoxia and chronic kidney disease by promoting oxygen delivery and carnitine homeostasis. Cell Metab. 2022;34(2):299–316.e6.

47. Bissinger R, Nemkov T, D’Alessandro A, et al. Proteinuric chronic kidney disease is associated with altered red blood cell lifespan, deformability and metabolism. Kidney Int. 2021;100(6):1227–1239.

48. D’Alessandro A, Nouraie SM, Zhang Y, et al. In vivo evaluation of the effect of sickle cell hemoglobin S, C and therapeutic transfusion on erythrocyte metabolism and cardiorenal dysfunction. Am J Hematol. 2023;98(7):1017–1028.

49. Wu H, Bogdanov M, Zhang Y, et al. Hypoxia-mediated impaired erythrocyte Lands’ Cycle is pathogenic for sickle cell disease. Scientific Reports 2016 6:1. 2016;6(1):1–16.

