## Supplemental Material for "Complete absence of GLUT1 does not impair human terminal erythroid differentiation"

#### Supplemental Material and Methods

##### Table S1 - Primary and secondary antibodies/RBD constructs used for flow cytometry

| Antibody name | Protein target | Species | Source | Dilution |
|---|---|---|---|---|
| anti-ASCT1.RBD.mFc | Neutral aminoacid transporter 1 | Mouse | Metafora | 1:20 |
| anti-ASCT2.RBD.mFc | Glutamine importer | Mouse | | 1:40 |
| anti-CAT1.RBD.rFc | Arginine importer 1 | Rabbit | | 1:20 |
| anti-FLVCR1.RBD.rFc | Heme Exporter | Rabbit | | 1:10 |
| anti-Glut1.RBD.mFc | Glucose transporter 1 | Mouse | | 1:5 |
| anti-MCT1.RBD.rFc | Lactate transporter 1 | Rabbit | | 1:5 |
| anti-PiT2.RBD.rFc | Phosphate importer 2 | Mouse | | 1:25 |
| anti-RFVT1/2.RBD.rFc | Riboflavin importers | Rabbit | | 1:20 |
| anti-SMVT.RBD.mFc | Na+ dependent multivitamin transporter | Mouse | | 1:5 |
| anti-XPR1.RBD.rFc | Phosphate Exporter | Rabbit | | 1:50 |
| BRIC 256 | GPA | Mouse IgG1 | IBGRL Research Products | 1:2 |
| BRIC216 | CD55 | Mouse IgG1 | | |
| BRIC221 | BCAM | Mouse IgG2b | | |
| BRIC222 | CD44 | Mouse IgG1 | | |
| BRIC32 | CD47 | Mouse IgG1 | | |
| BRIC4 | GPC | Mouse IgG1 | | |
| BRIC68 | KELL | Mouse IgG2a | | |
| BRIC69 | Rh | Mouse IgG1 | | |
| BRIC71 | Band 3 | Mouse IgG1 | | |
| LA1818 | RhAG | Mouse IgG1 | | |
| Purified anti-human CD147 (HIM6) | Basagin | Mouse IgG1 | Biolegend | 1:50 |
| Purified Mouse IgG1 (MG1-45) | Mouse IgG1 | Mouse IgG1 | | |
| Purified Mouse IgG2a (MOPC-173) | Mouse IgG2a | Mouse IgG2a | | |
| Purified Mouse IgG2b (MPC-11) | Mouse IgG2b | Mouse IgG2b | | |

| | | | | |
|---|---|---|---|---|
| APC anti-human CD71 (CY1G4) | Transferrin receptor | Rat IgG2a | | |
| APC anti-mouse IgG1 (RMG1-1) | Mouse IgG1 | Mouse IgG1 | Biolegend | |
| APC Goat anti-mouse IgG (Poly4053) | Mouse IgG | Goat Polyclonal IgG | | |
| FITC anti-Glut1.RBD | Glucose transporter 1 | Mouse | Metafora | 1:50 |
| FITC anti-human CD49d (MZ18-24A9) | CD49d | Mouse IgG2b | Miltenyi Biotech | |
| FITC Isotype control, mouse IgG2b (IS6-11E5.11 | Mouse IgG2b | Mouse IgG2b | | |

**Table S2 - Antibodies and reagents used for immunoblotting.**

| Antibody name | Protein target | Species | Source | Dilution |
|---|---|---|---|---|
| sc-25731 | Adducin-alpha | Mouse | Santa Cruz Biotechnology | 1:1000 |
| sc-47724 | GAPDH | Mouse | | 1:2000 |
| anti-Glut1 | Glucose transporter 1 | Rabbit | in-house product | 1:5000 |
| anti-stomatin | Stomatin | Rabbit | | 1:1000 |
| Rabbit Anti-mouse (P0260) HRP | Mouse IgG | Rabbit Polyclonal | Dako | 1:2000 |
| Swine Anti-Rabbit (P0399) HRP | Rabbit IgG | Swine Polyclonal | | |

Chemiluminescence detection was done using Cytivas ECL WB detection reagents (RNP2106).

#### CD34+ culture

Peripheral blood mononuclear cells are isolated from waste apheresis cones[19], followed by CD34+ magnetic cell isolation with CD34+ MicroBead kit according to manufacturer's protocol. Cells were cultured as described by Kupzig et al [20]. Isolated cells were seeded at 1-2 x10$^5$/mL in IMDM base medium. From days 0 to 8 (1$^{st}$ stage) medium is supplemented with 40ng/mL SCF and 1ng/mL IL-3, and from days 8-13 (2$^{nd}$ stage) only with 40ng/mL SCF. From day 13 onwards (3$^{rd}$ stage) no supplementation is required. Cells were kept in culture until days 20 or 21. To obtain a purified population of reticulocytes, the cultures were filtered by leukofiltration. A leukocyte reduction filter (Macopharma) was pre-soaked and equilibrated with phosphate-buffered saline (PBS) and the cultured cell suspension was loaded into the filter followed by at least three volumes of PBS and allowed to pass through under gravity. The resulting flow-through was then centrifuged at 400 × g, for 15 min and the pelleted cells were resuspended in PBSAG (PBS + 1 mg/ml BSA, 2 mg/ml glucose) and kept at 4 °C.

#### Single clone selection of BEL-A GLUT1 KO

To obtain BEL-A GLUT1 clonal cell lines, 72h after nucleofection, the mixed population was sorted into single clones and genomic DNA (gDNA) was extracted using QuickExtract (Lucigen) following supplier's protocol. The region of interest was amplified through polymerase chain reaction (PCR) using Q5 High-Fidelity 2X Master Mix (NEB) and the primers SLC2A1 Forward 5'-CAACCCCTTGTTTCTCGCCTGG and Reverse 5'- AAGGGCTGTGGGTGACACTTCA. The samples were purified using the QIAquick PCR Purification Kit (Qiagen) before being sent to

Eurofins for sequencing. Clones with mutations resulting in frameshift mutations with predicted loss-of-expression were labelled for GLUT1, confirming through flow cytometry the sequencing results.

##### *P. falciparum* invasion assays

*P. falciparum* strain 3D7 parasites (BEI Resources) were maintained in human erythrocytes at 5% hematocrit using standard culture conditions[19]. Schizont stage parasites were magnetically purified using the Magnetic Cell Separation (MACS) system (Miltenyi Biotec) and added to wells of a round bottomed 96-well plate containing $1x10^6$ leukofiltered CD34-derived reticulocytes. Parasitemias ranging from 1-8% were used. Heparin (100 mU/µl final) was used to inhibit invasion in negative controls. After ~18 h, invasion was quantified using flow cytometry. For flow cytometry, cells were stained with SYBR Green (1:2000 in culture media; Sigma-Aldrich) for 30 min at 37°C in the dark. Cells were centrifuged, SYBR Green containing media removed and then fixed for 15 minutes at room temperature as described above. Invasion was quantified based on SYBR green positivity using the heparin control to correct for background events within the invasion gate.

##### Reverse transcription quantitative-PCR (RT-qPCR)

RNA was isolated from frozen pellets of expanding BEL-A cells using a Rneasy kit (Qiagen), following supplier's protocol. Total RNA concentration was measured using NanoDrop (ThermoFisher). Complementary DNA (cDNA) was generated using the QuantiTect Reverse Transcription Kit (Qiagen) following the indicated protocol. The RT-qPCR was done in 20µL reactions in a MicroAmpTM Optical 96-Well Reaction Plate (Applied Biosystems). Assuming a total conversion of RNA to cDNA, 10ng of cDNA was used as template with 10µL of PowerUpTM SYBRTM Green Master Mix (Applied Biosystems), 1µL of each primer (10µM) and 6µL nuclease-free water. Each sample was analysed in triplicate and a no template control was included. The RT-qPCR were performed using the QuantiStudio3™ RT PCR system (Applied Biosystems) under the Standard cycling mode (Primer $T_m$ > 60°C) as indicated by the manufacture's protocol for SYBR green dyes. The relative gene expression was determined with the $2^{-\Delta\Delta Ct}$ method. Primers (Eurofins) used were GLUT1 5'- GAGAACCGGGCCAAGAGTGTGC and 5'-AGCGGAACAGCTCCAGGATGGT; GLUT3 5'-TCGGCTCCTTTTCCGTCGGACT and 5'-AAAGCAGCCACCAGTGACAGCC; GAPDH 5'- GAGTCAACGGATTTGGTCGT and 5'-TTGATTTTGGAGGGATCTCG.

##### Metabolomics

Metabolomics analyses were performed as previously described[23]. Cells were extracted in ice cold 5:3:2 MeOH:MeCN:water (v/v/v) at a $1x10^6$ cells/ml ratio, then vortexed for 30 min at 4 °C. Supernatants were clarified by centrifugation (10 min, 12,000 g, 4 °C). The resulting metabolite extracts were analysed (10 uL per injection) by ultra-high-pressure liquid chromatography coupled to mass spectrometry (UHPLC-MS — Vanquish and QExactive, Thermo). Metabolites were resolved on a Phenomenex Kinetex C18 column (2.1 x 150 mm, 1.7 um) at 45 °C using a 5-minute gradient method in positive and negative ion modes (separate runs) over the scan range 65-975 m/z exactly as previously described[23]. Oxylipins were resolved on a Waters ACQUITY UPLC BEH C18 column (2.1 x 100 mm, 1.7 µm) at 60 °C using mobile phase (A) of 20:80:0.02 MeCN:water:formic acid (FA) and a mobile phase (B) of 20:80:0.02 MeCN:isopropanol:FA. For negative mode analysis the chromatographic the gradient was as follows: 0.35 mL/min flowrate, 0% B 0-0.5 min, 25% B at 1 min, 40% B at 2.5min, 55% B at 2.6min, 70% B at 4.5 min, 100% B at 4.6-6 min, 0% B at 6.1-7 min. The Q Exactive MS was operated in negative ion mode, scanning in Full MS mode (2 µscans) from 150 to 1500 m/z at 70,000 resolution, with 4 kV spray voltage, 45 sheath gas, 15 auxiliary gas. Following data acquisition, .raw files were converted to .mzXML using RawConverter then metabolites assigned and peaks integrated using Maven (Princeton University) in conjunction with the KEGG database and an in-house standard library. Quality control was assessed as using technical replicates run at beginning, end, and middle of each sequence as previously described[24,25].

#### Lipidomics

Total lipids were extracted as previously described[26]: cells were extracted in cold methanol at a $1x10^6$ cells/ml ratio. Samples were then briefly vortexed and incubated at -20 °C for 30 minutes. Following incubation, samples were centrifuged at 12,700 RPM for 10 minutes at 4 °C and 80 µL of supernatant was transferred to a new tube for analysis. Lipid extracts were analyzed (10 uL per injection) on a Thermo Vanquish UHPLC/Q Exactive MS system using a 5 min lipidomics gradient and a Kinetex C18 column (30 x 2.1 mm, 1.7 µm, Phenomenex) held at 50 °C. Mobile phase A: 25:75 MeCN:water with 5 mM ammonium acetate; Mobile phase B: 90:10 isopropanol:MeCN with 5 mM ammonium acetate. The gradient and flow rate were as follows: 0.3 mL/min of 10% B at 0 min, 0.3 mL/min of 95% B at 3 min, 0.3 mL/min of 95% B at 4.2 min, 0.45 mL/min 10% B at 4.3 min, 0.4 mL/min of 10% B at 4.9 min, and 0.3 mL/min of 10% B at 5 min. Samples were run in positive and negative ion modes (both ESI, separate runs) at 125 to 1500 m/z and 70,000 resolution, 4 kV spray voltage, 45 sheath gas, 25 auxiliary gas. The MS was run in data-dependent acquisition mode ($ddMS^2$) with top10 fragmentation. Raw MS data files were searched using LipidSearch v 5.0 (Thermo).

#### Proteomics

Proteomics analyses were performed as described[27]. A volume of 10 µL of cells were lysed in 90 µL of distilled water; 5 µL of lysates were mixed with 45 µL of 5% SDS and then vortexed. Samples were reduced with 10 mM DTT at 55 °C for 30 min, cooled to room temperature, and then alkylated with 25 mM iodoacetamide in the dark for 30 min. Next, a final concentration of 1.2% phosphoric acid and then six volumes of binding buffer (90% methanol; 100 mM triethylammonium bicarbonate, TEAB; pH 7.1) were added to each sample. After gentle mixing, the protein solution was loaded to a S-Trap 96-well plate, spun at 1500 x g for 2 min, and the flow-through collected and reloaded onto the 96-well plate. This step was repeated three times, and then the 96-well plate was washed with 200 µL of binding buffer 3 times. Finally, 1 µg of sequencing-grade trypsin (Promega) and 125 µL of digestion buffer (50 mM TEAB) were added onto the filter and digested carried out at 37 °C for 6 h. To elute peptides, three stepwise buffers were applied, with 100 µL of each with one more repeat, including 50 mM TEAB, 0.2% formic acid (FA), and 50% acetonitrile and 0.2% FA. The peptide solutions were pooled, lyophilized, and resuspended in 500 µL of 0.1 % FA.

Each sample was loaded onto individual Evotips for desalting and then washed with 200 µL 0.1% FA followed by the addition of 100 µL storage solvent (0.1% FA) to keep the Evotips wet until analysis. The Evosep One system (Evosep, Odense, Denmark) was used to separate peptides on a Pepsep column, (150 um inter diameter, 15 cm) packed with ReproSil C18 1.9 um, 120A resin. The system was coupled to a timsTOF Pro mass spectrometer (Bruker Daltonics, Bremen, Germany) via a nano-electrospray ion source (Captive Spray, Bruker Daltonics). The mass spectrometer was operated in PASEF mode. The ramp time was set to 100 ms and 10 PASEF MS/MS scans per topN acquisition cycle were acquired. MS and MS/MS spectra were recorded from *m/z* 100 to 1700. The ion mobility was scanned from 0.7 to 1.50 Vs/cm2. Precursors for data-dependent acquisition were isolated within ± 1 Th and fragmented with an ion mobility-dependent collision energy, which was linearly increased from 20 to 59 eV in positive mode. Low-abundance precursor ions with an intensity above a threshold of 500 counts but below a target value of 20000 counts were repeatedly scheduled and otherwise dynamically excluded for 0.4 min.

#### Database Searching and Protein Identification

MS/MS spectra were extracted from raw data files and converted into .mgf files using MS Convert (ProteoWizard, v. 3.0). Peptide spectral matching was performed with Mascot (v. 2.5) against the Uniprot human database. Mass tolerances were +/- 15 ppm for parent ions, and +/- 0.4 Da for fragment ions. Trypsin specificity was used, allowing for 1 missed cleavage. Protein N-terminal acetylation, isopeptide bond formation with loss of ammonia (K), and peptide N-terminal pyroglutamic acid formation were set as variable modifications with Cys carbamidomethylation set as a fixed modification.
Scaffold (v 4.8, Proteome Software, Portland, OR, USA) was used to validate MS/MS based peptide and protein identifications. Peptide identifications were accepted if they could be established at greater than 95.0% probability as specified by the Peptide Prophet algorithm. Protein identifications were accepted if they could be established at greater than 99.0% probability and contained at least two identified unique peptides.

#### Statistics

The non-parametric Mann-Whitney U test or Kruskal-Wallis test with Bonferroni correction were used to test for differences between groups. *$p < 0.05$, **$p < 0.01$, ***$p < 0.001$
For omics data, statistical analysis were conducted using MetaboAnalyst v 5.0 and Rstudio upon autoscale normalization (i.e., data were mean-centered, divided by the standard deviation of each variable). Line plots and volcano plots of correlations (Spearman) were generated upon analysis of the raw data via Rstudio. Significance was calculated upon false discovery rate correction. Network analyses and pathway analyses were performed in OmicsNet v 2.0 using as input the significantly altered metabolites and proteins (FDR-corrected $p < 0.05$).

### Supplemental Figures

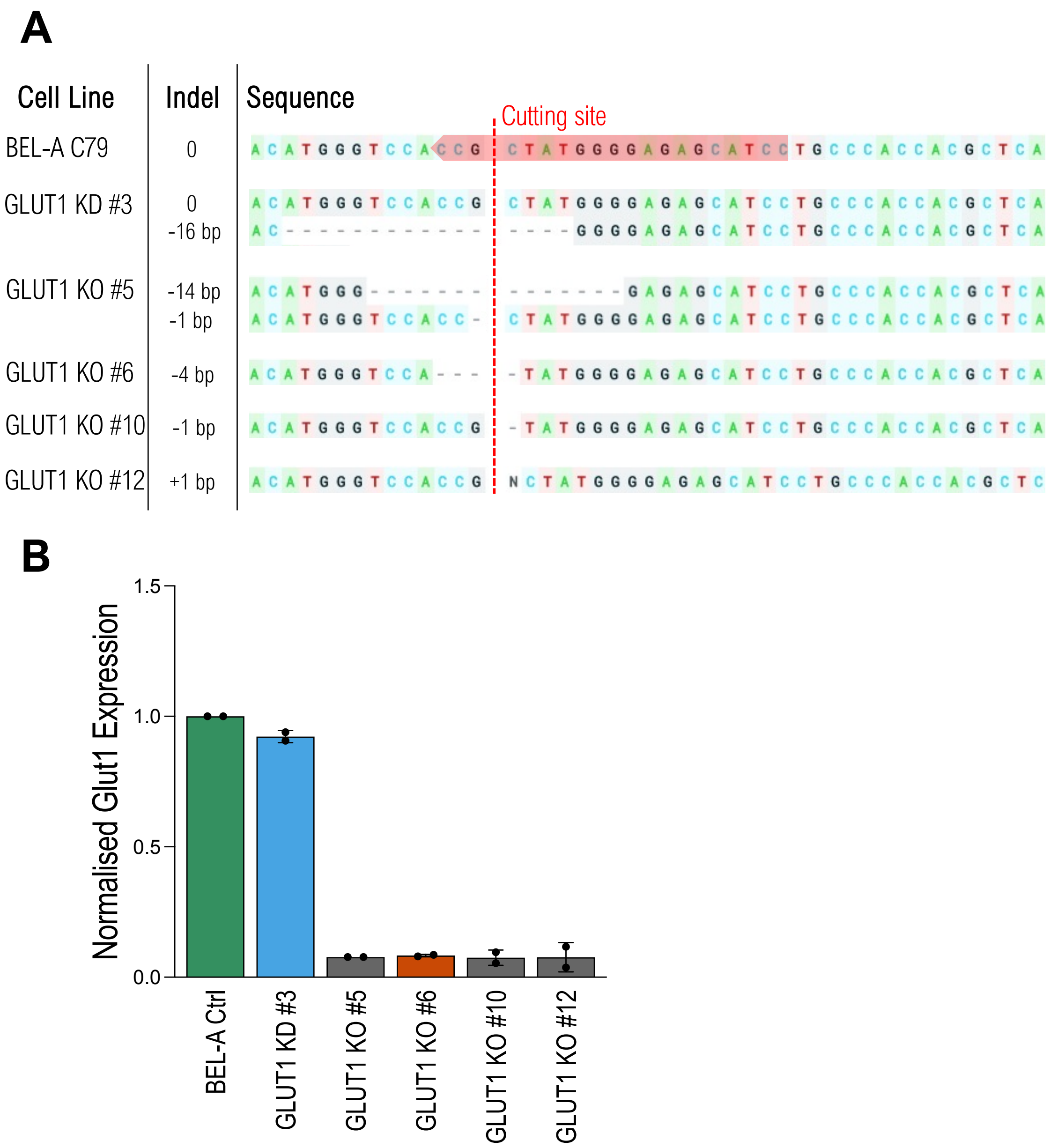


**Supplemental Figure 1 – GLUT1 gene editing of BEL-A cell line using CRISPR-Cas9 . GLUT1 KD 3 and KO 6 are mentioned in the main manuscript**

A Sequencing of SLC2A1 (GLUT1) gene on BEL-A C79, highlighting the gRNA (red) used for CRISPR editing. Analysis was performed using Synthego's ICE tool. Sanger sequencing of clonal edited lines uncovered clone 3 as a knockdown and clones 5, 6, 10, and 12 as knockdowns. All mutations shown are in the vicinity of the cutting site (red line). B Bar graph illustrating GLUT1 expression on expanding BEL-A cell lines. The median fluorescence intensity is normalised for endogenous expression of BEL-A C79. Results are means ± standard deviation (SD), n=2.

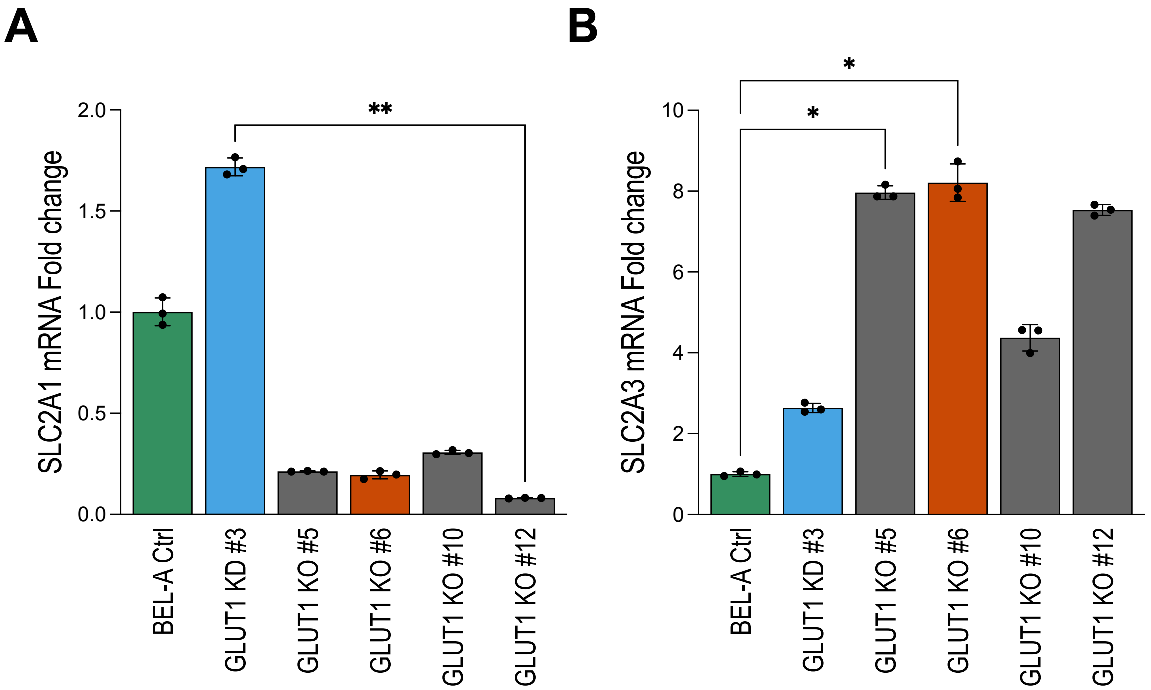


**Supplemental Figure 2: mRNA expression of SLC2A1 and SLC2A3 measured by real-time quantitative PCR on wildtype and GLUT1-mutated BEL-A cell lines**

Analysis of GLUT1 (**A**), and GLUT3 (**B**) mRNA levels in expanding BEL-A GLUT1 KD/KO clonal cell lines by real time quantitative PCR (RT-qPCR). Target gene expression was normalised to GAPDH reference gene and to BEL-A C79 expression, using the 2-ΔΔCt method. Results are means ± SD, n=3. Significance was assessed with the non-parametric Kruskal-Wallis test with Dunn's correction, *p<0.05, **p<0.01

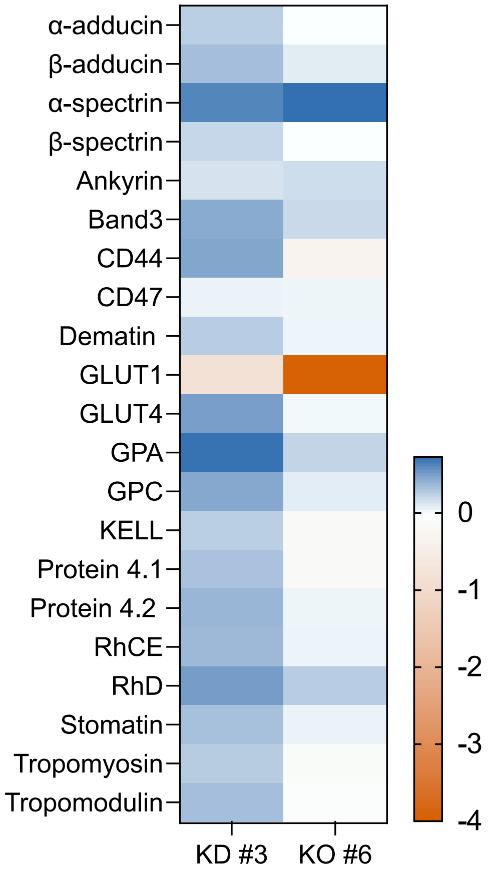


**Supplemental Figure 3 - Proteomics confirm GLUT1 KO in BEL-A cells and show no impact in main membrane or cytoskeletal proteins**

Heat map visualisation of the tandem mass tag (TMT) proteome subset generated from $\log_2$ fold-change values of membrane and cytoskeletal proteins expression in paired samples of GLUT1 KD #3 and KO #6 with BEL-A C79 control cell line as baseline. Red denotes low expression and dark green high expression.

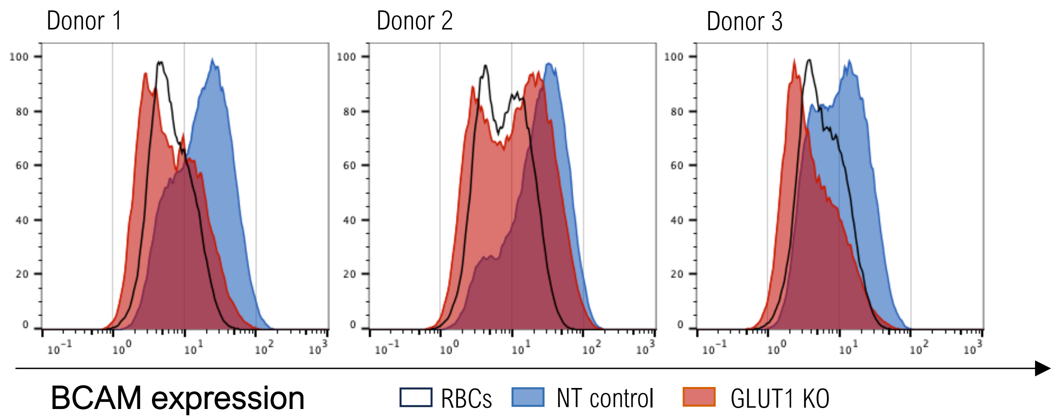


**Supplemental Figure 4 – Basal cell adhesion molecule (BCAM) expression is altered in GLUT1-KO primary reticulocytes**

Flow cytometry histograms of BCAM expression on NT-controls (green) and GLUT1-KO (red) cultured reticulocytes, including the matched RBCs expression (black line) of three individual donors.

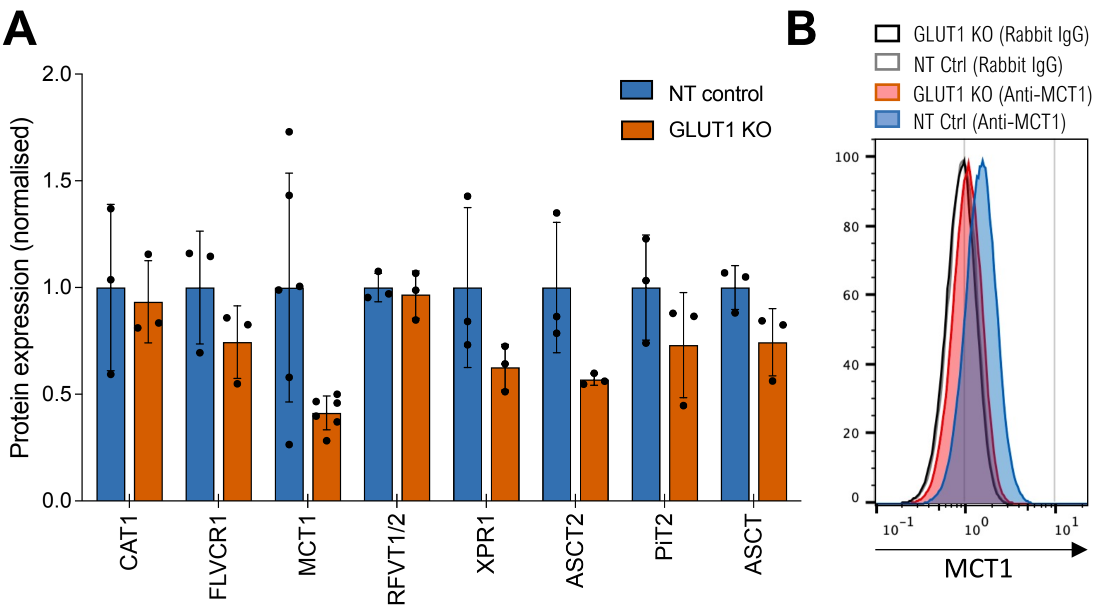


**Supplemental Figure 5 – Membrane expression of various nutrient transporters detected used RBD reagents**

A Bar plots of nutrient transporters expression assessed with RBD constructs normalised to NT controls. Staining done on fixed CD34-derived reticulocytes. (n=3, SD, MCT1 has 2 technical repeats per donor). Arginine Importer (CAT1), Heme exporter (FLVCR1), Lactate transporter 1 (MCT1), Riboflavin importers (RFVT1/2), Phosphate Exporter (XPR1), Glutamine Importer (ASCT2), Phosphate Importer 2 (PiT2), and Neutral Aminoacid Transporter 1 (ASCT1). B Histogram exemplifying MCT1 labelling on filtered CD34-derived reticulocytes from one donor.

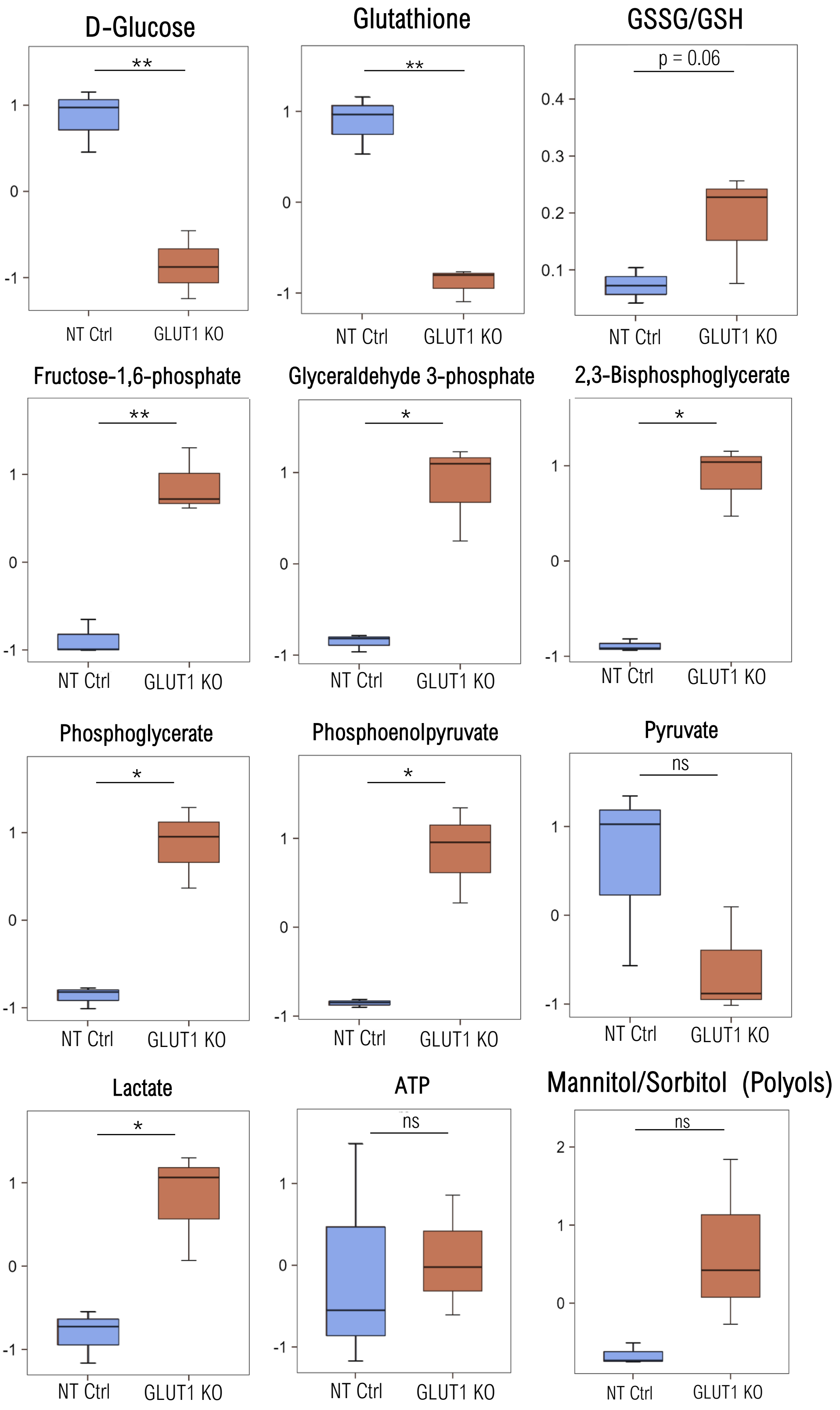


**Supplemental Figure 6 – Metabolite analysis between NT-control and GLUT1-KO primary reticulocytes**

Box plots analysis (mean±min to max with standard deviation) displayed as Log2 (fold change). Data analysis performed by RStudio, and significance was calculated upon false discovery rate correction ($p < 0.05$).

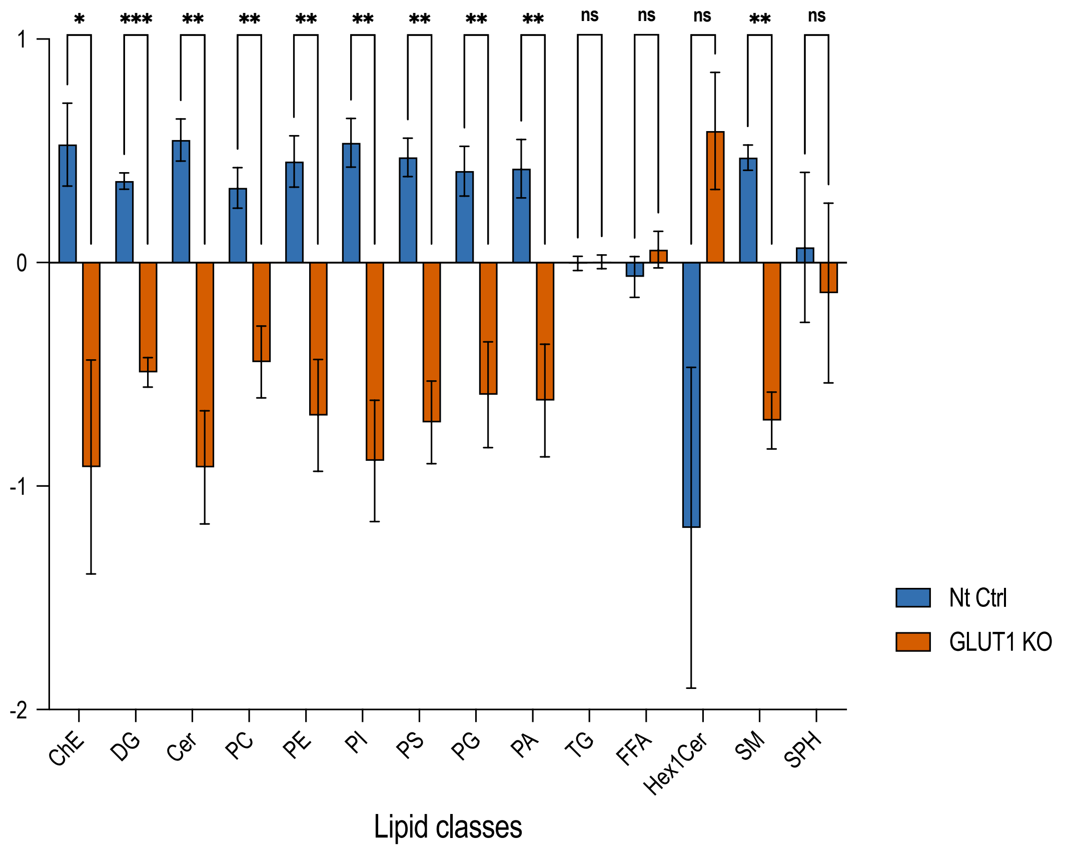


**Supplemental Figure 7 – Lipid class analysis between NT-control and GLUT1-KO primary reticulocytes**

Lipidomics results comparing lipid classes between NT-control and GLUT1-KO, indicating the levels of cholesteryl-esters (ChE), diacylglycerols (DG), ceramides (Cer), all phospholipids (PC, PE, PI, PS, PG, PA), triacylglycerols (TG), free fatty acids (FFA), mono-hexosylceramides (Hex1Cer)., sphingomyelin (SM), and sphingophospholipids (SPH). Data shown in the fold change (log2) of the sum of all lipid chains of each class normalised for the median of of 6 samples (n=3, for each group). Significance was assessed by multiple t-tests, *$p<0.05$, **$p<0.01$, and ***$p<0.001$.
